# Tobacco smoke exposure results in recruitment of inflammatory airspace monocytes and accelerated growth of *Mycobacterium tuberculosis*

**DOI:** 10.1101/2022.12.21.521304

**Authors:** Bjӧrn Corleis, Constantine N. Tzouanas, Marc H Wadsworth, Josalyn L Cho, Alice H Linder, Abigail E Schiff, Amy K Dickey, Benjamin D Medoff, Alex K. Shalek, Douglas S Kwon

**Affiliations:** Ragon Institute of MGH, MIT, and Harvard; Cambridge, MA USA; Institute of Immunology, Friedrich-Loeffler-Institute; Greifswald-Insel Riems, Germany; Institute for Medical Engineering & Science (IMES), Department of Chemistry, and Koch Institute for Integrative Cancer Research, MIT; Cambridge, Massachusetts, USA; Broad Institute of MIT and Harvard; Cambridge, Massachusetts, USA; University of Iowa Roy J and Lucille A Carver College of Medicine, Department of Internal Medicine, Division of Pulmonary, Critical Care and Occupational Medicine; Iowa City, Iowa, United States; Department of Medicine, Brigham and Women’s Hospital, Boston, MA, USA; Division of Pulmonary and Critical Care Medicine, Massachusetts General Hospital; Boston, MA USA; Department of Medicine, Harvard Medical School; Boston, MA, USA; Department of Immunology, Harvard Medical School, Boston; MA, USA; Division of Infectious Diseases, Massachusetts General Hospital; Boston, MA, USA

## Abstract

Tobacco smoking doubles the risk of active tuberculosis (TB) and accounts for up to 20% of all active TB cases globally. How smoking promotes lung microenvironments permissive to *Mycobacterium tuberculosis* (*Mtb*) growth remains incompletely understood. We investigated primary bronchoalveolar lavage cells from current- and never-smokers by performing single-cell RNA-sequencing (scRNA-seq), flow cytometry, and functional assays. We observed enrichment of immature inflammatory monocytes in the lungs of smokers compared to non-smokers. These monocytes exhibited phenotypes consistent with recent recruitment from blood, ongoing differentiation, increased activation, and states similar to those with chronic obstructive pulmonary disease (COPD). Using integrative scRNA-seq and flow cytometry, we identify CD93 as a marker for a subset of these newly recruited smoking-associated lung monocytes and further provide evidence that recruitment of monocytes into the lung is mediated by CCL11 binding to CCR2. We also show that these cells exhibit elevated inflammatory responses upon exposure to *Mtb* and accelerated intracellular growth of *Mtb* compared to mature macrophages. This elevated *Mtb* growth could be inhibited with an anti-inflammatory small molecule, providing a direct connection between smoking-induced pro-inflammatory states and permissiveness to *Mtb* growth. Our findings suggest a model in which smoking leads to recruitment of immature inflammatory monocytes from the periphery to the lung via CCL11-CCR2 interactions, which results in the accumulation of these *Mtb* permissive cells in the airway. This work defines how smoking may lead to increased susceptibility to *Mtb* and identifies novel host-directed therapies to reduce the burden of TB among those who smoke.

**One Sentence Summary:** Inflammatory monocytes are recruited to the airways of smokers where they may contribute to more rapid growth of *Mycobacterium tuberculosis* in the lungs.

## INTRODUCTION

Tuberculosis (TB) has infected over 20% of the global population, and active TB kills 1-2 million people every year (*1*). Current treatment regimens are poorly effective and require administration of multiple antibiotics for months, highlighting the need for greater mechanistic understanding of disease to inform novel preventative and therapeutic strategies (*2*). The majority of people infected with the causative agent of TB, *Mycobacterium tuberculosis* (*Mtb*), develop subclinical latent TB infection (LTBI), with only 5-10% of individuals with LTBI eventually developing clinical disease (*3*). While several risk factors are linked to active TB and TB mortality, one of the strongest is smoking (*1*). Globally, 1.3 billion people smoke tobacco, which results in an estimated ten to twenty percent of all annual active TB cases being attributed to smoking (*4, 5*). Despite the tremendous contribution of smoking to the worldwide burden of TB, very little is understood about the specific mechanisms by which smoking promotes active TB.

Upon entry, *Mtb* primarily proliferates within mature alveolar macrophages (AMs). However, the phenotype and metabolic profiles of AMs from tobacco smokers are significantly altered and associated with diminished phagocytosis and anti-mycobacterial defense mechanisms (*4, 6–11*). Furthermore, tobacco smoking has been reported to change the cellular composition of other immune lineages in this compartment. For instance, we recently reported a relative decrease of the lymphocyte population and an increase in the total number of macrophages in the airspace compartment with smoking (*9, 12*). However, a comprehensive understanding of which cell types and states are altered with smoking and how these potentially impact the risk of active TB is lacking. A more detailed understanding of how smoking perturbs the composition and state of the lung microenvironment would provide insights into the mechanisms underlying smoking-associated increased risk for active TB and disease progression, as well as potential avenues for the development of host-directed therapies to prevent and treat active TB.

In this study, we demonstrate that tobacco smoking leads to the recruitment of immature inflammatory monocytes to the lung. These recruited monocytes exhibit phenotypic and transcriptional hallmarks of elevated inflammatory responses and the potential to differentiate into monocyte-derived macrophages (MDMs). We identify CD93 as a marker for a subset of newly recruited airspace monocytes and further provide *ex vivo* evidence that recruitment of monocytes into the lung is mediated by CCL11 binding to CCR2. We demonstrate that blood and lung monocytes from smokers exhibit elevated inflammatory responses following exposure to *Mtb* and that immature monocytes support increased rates of *Mtb* growth relative to mature macrophages. Further, we show that treatment of these monocytes with anti-inflammatory drugs can inhibit elevated intracellular *Mtb* growth. These findings provide new biological understanding of the mechanism by which smoking increases *Mtb* growth and reveal potential strategies to mitigate TB burden among tobacco smokers.

## RESULTS

### Tobacco smoking leads to an increase of monocytes in the air spaces

To explore the effects of smoking on airspace cells in the lung, we characterized broncho-alveolar-lavage (BAL) samples from smokers and non-smokers using scRNA-seq, flow cytometry, and *ex vivo* functional assays (Figure 1A). We found that the total cellularity of BAL fluid was significantly increased in smokers compared to non-smokers (Figure S1A). We identified airspace monocytes by flow cytometry, which were distinct from alveolar macrophages (AMs) in their size, granularity, and surface expression of CD14 (Figure S1B-C). There was a 7-fold increase in the number of small (SSC/FSC^low^) CD14^+^ airspace monocytes in smokers versus non-smokers, with no significant difference in AMs (SSC/FSC^high^CD45^+^CD14^low^CD16^+^), granulocytes (PMN), and T cells (Figure 1B, S1B-C). Thus, our data indicate that the increase in total airspace cells was primarily driven by an increase in small CD14^+^ airspace monocytes.

**Fig. 1.**
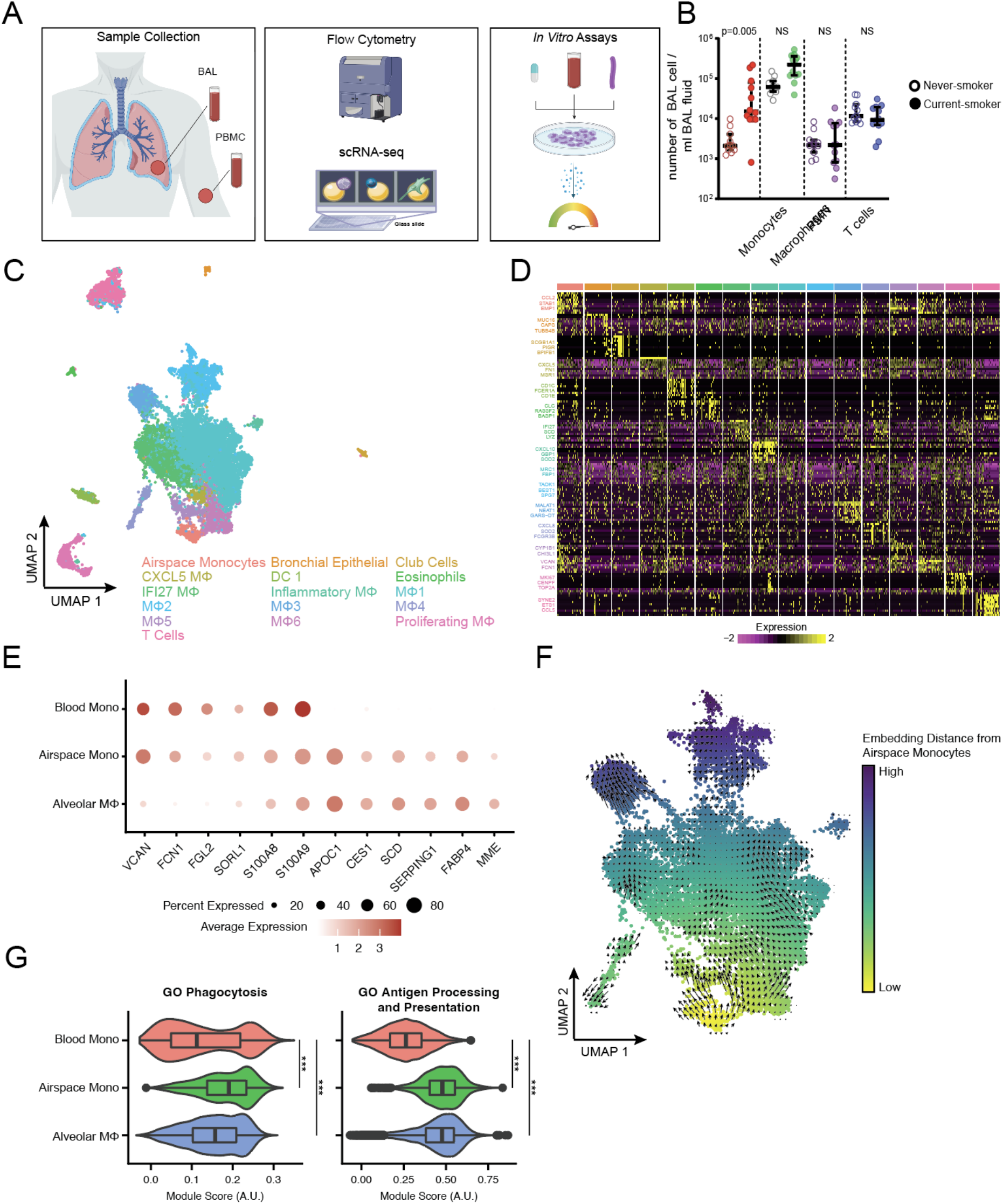
Increased number of differentiating airspace monocytes in the lung of tobacco smokers. (A) Schematic of the sample processing pipeline using Seq-Well S^3^, flow cytometry and *in vitro* assays. (B) Analysis of total number of viable cells per ml BAL fluid in non-smokers versus smokers showed significantly higher number of airspace monocytes in smokers. (C) Uniform Manifold Approximation and Project (UMAP) plot (n = 20,799 cells, N = 9 individuals) colored by cell types. (D) Heatmap showing normalized gene expression values (log (scaled UMI + 1)) for cell-type-defining genes. (E) Dotplot of myeloid-associated marker genes that distinguish blood monocytes and macrophages, with airspace monocytes exhibiting an intermediate phenotype. (F) RNA velocity analysis of differentiation trajectories among myeloid cells derived from BAL samples, colored by distance in embedding space from airspace monocyte cluster. (G) Module scoring for GO processes relevant to monocyte/macrophage function. P < 0.001 by one-way ANOVA for each GO process, *** indicates Benjamini-Hochberg-corrected p-value < 10^-10^. Box-and-whiskers show median and interquartile range.

To further characterize the cells across the lung and blood compartments, we performed picowell-based scRNA-seq on BAL and PBMCs from smokers (n=5) and never-smokers (n=4). After performing initial quality controls, we retained 20,799 BAL cells and 36,405 PBMCs for further analysis (Figure S2). Dimensionality reduction, unsupervised clustering, and manual annotation identified 16 populations among the BAL samples, with myeloid cells as the predominant cell type (Figure 1C-D, Figure S2A-C) (*13, 14*). We chose to prioritize airspace monocytes for further investigation, as they were significantly enriched in the lungs of smokers as measured through both flow cytometry (Figure 1B; p = 0.005, Wilcoxon rank sum test) and scRNA-seq (Figure S2D; p = 0.0049, Dirichlet regression analysis, which accounts for compositional shifts in one cell type necessarily affecting relative proportions of other cell types). Expression profiles from the airspace monocytes best aligned with monocytes in a previously published blood immune atlas (*15*) (Figure S3A) and exhibited expression of myeloid-associated markers at levels intermediate between blood-derived monocytes and AMs (Figure 1E). To understand the differentiation potential of airspace monocytes, we conducted RNA velocity analyses, which leverage unspliced-to-spliced mRNA ratios to infer cells’ dynamic trajectories from scRNA-seq readouts (*16, 17*). In this approach, a gene with a high unspliced-to-spliced mRNA ratio would indicate recent transcriptional activation and potential information about a cell’s future state. RNA velocity analyses indicated that airspace monocytes are an immature cell type with the capacity to differentiate into more mature macrophage populations, with trajectories originating in the airspace monocyte cluster and moving towards more mature macrophage cell clusters (i.e., Macrophage 5 and Macrophage 6; Figure 1F). Consistent with the notion that these airspace monocytes are an infiltrating, immature population associated with smoking, airspace monocytes, Macrophage 5, and Macrophage 6 populations (i.e., the macrophage clusters predicted to be descendants of airspace monocytes) were also enriched in samples from smokers compared to never-smokers (Fig. S2D-E; Macrophage 5 p-value = 6×10^-5^; Macrophage 6 p-value = 7.3×10^-6^; Dirichlet regression analysis). Both airspace monocytes and macrophages exhibited elevated expression of genes involved in phagocytosis and antigen processing/presentation as compared to blood monocytes from the same subjects (Figure 1G and Figure S3B), supporting the interpretation that airspace monocytes are a differentiating cell population. As external support from a separate human disease cohort, we compared our airspace monocyte population against work from Baβler et al. (*18*), who identified a population of monocytes differentiating into macrophages that was enriched in BAL samples from smokers with chronic obstructive pulmonary disease (COPD). Our smoking-associated airspace monocytes strongly and specifically expressed markers from a population of differentiating monocytes (i.e., elevated compared to blood-derived monocytes and AM alike) identified in Baβler et al., further supporting that airspace monocytes accumulate in smokers and may serve as precursors to other tissue macrophage states (Figure S3C-E). This prior work likewise found that COPD-linked differentiating monocytes exhibited associations with smoking and elevated expression of lipid metabolism genes, both of which were concordant with our dataset and reflected through compositional analyses (Figure 1B for flow cytometry compositional analyses; Figure S2D-E for scRNA-seq compositional analyses), gene expression analyses (Figure S3C-E), and histological staining of BAL cells for neutral lipids (Figure S3F-G). Importantly, phenotypic gene modules linked to these disease-associated monocytes (i.e., infiltration/differentiation, lipid dysregulation) were upregulated in airspace monocytes from smokers compared to those from never-smokers in our dataset, further supporting the link between smoking and perturbed monocyte states (Figure S3D). As a control, we confirmed that the differentiating monocyte phenotype was not correlated with potential cohort structure confounders such as age or biological sex but was correlated with other previously published smoking-related gene sets in airspace immune cells (Figure S3H) (*19–22*).

In summary, we used flow cytometry and scRNA-seq to identify an airspace monocyte population that is preferentially enriched in the airspaces of tobacco smokers and exhibits phenotypes intermediate between circulating blood monocytes and more mature macrophages in the airspaces. RNA velocity analyses supported these airspace monocytes as progenitors for other macrophage subpopulations that are also enriched in smokers, and comparison to prior studies revealed concordant upregulation of genes observed in disease-associated monocyte phenotypes.

### Airspace monocytes originate from blood monocytes and share the expression of CD93

To further characterize the airspace monocytes enriched in smokers, we compared gene signatures of airspaces versus blood monocytes from the same donors. Differential expression and gene set enrichment analysis revealed enrichment in airspace monocytes for pathways associated with immune activation, upregulation of defense mechanisms, and locomotion (Figure 2A-B and Figure S4) compared to blood monocytes. To better understand connections between airspace and blood monocytes, we used pseudotime analyses to find a trajectory that preserves the high-dimensional structure of our scRNA-seq dataset while ordering cells according to smooth gradients in gene expression (*23*). Complementing the orthogonal RNA velocity analyses (Figure 1F), these results suggested that airspace monocytes originate from blood monocytes and assume intermediate positions along the differentiation trajectory towards more mature macrophage states (Figure 2C). They also revealed genes that vary with smoking-associated recruitment, infiltration, and differentiation, such as: 1. genes elevated early in pseudotime (i.e., associated with blood monocytes) but rapidly lost (e.g., signaling receptors like *CSF3R*, *CX3CR1*, and *CXCR4*; inflammation-associated genes like *F13A1*, *DUSP1*, and *CTSS*); 2. genes that were elevated at intermediate stages of the pseudotime continuum from monocyte to macrophage (e.g., secreted proteins like *CTSL*, *C1QA*; cell surface-related genes like *CD81*, *DST*, and *DAB2*); and, 3. markers of mature macrophages that only increase at the terminus of the pseudotime ordering (e.g., *CD163*, *FCGRT*, *CD68*) (Figure 2D).

**Fig. 2.**
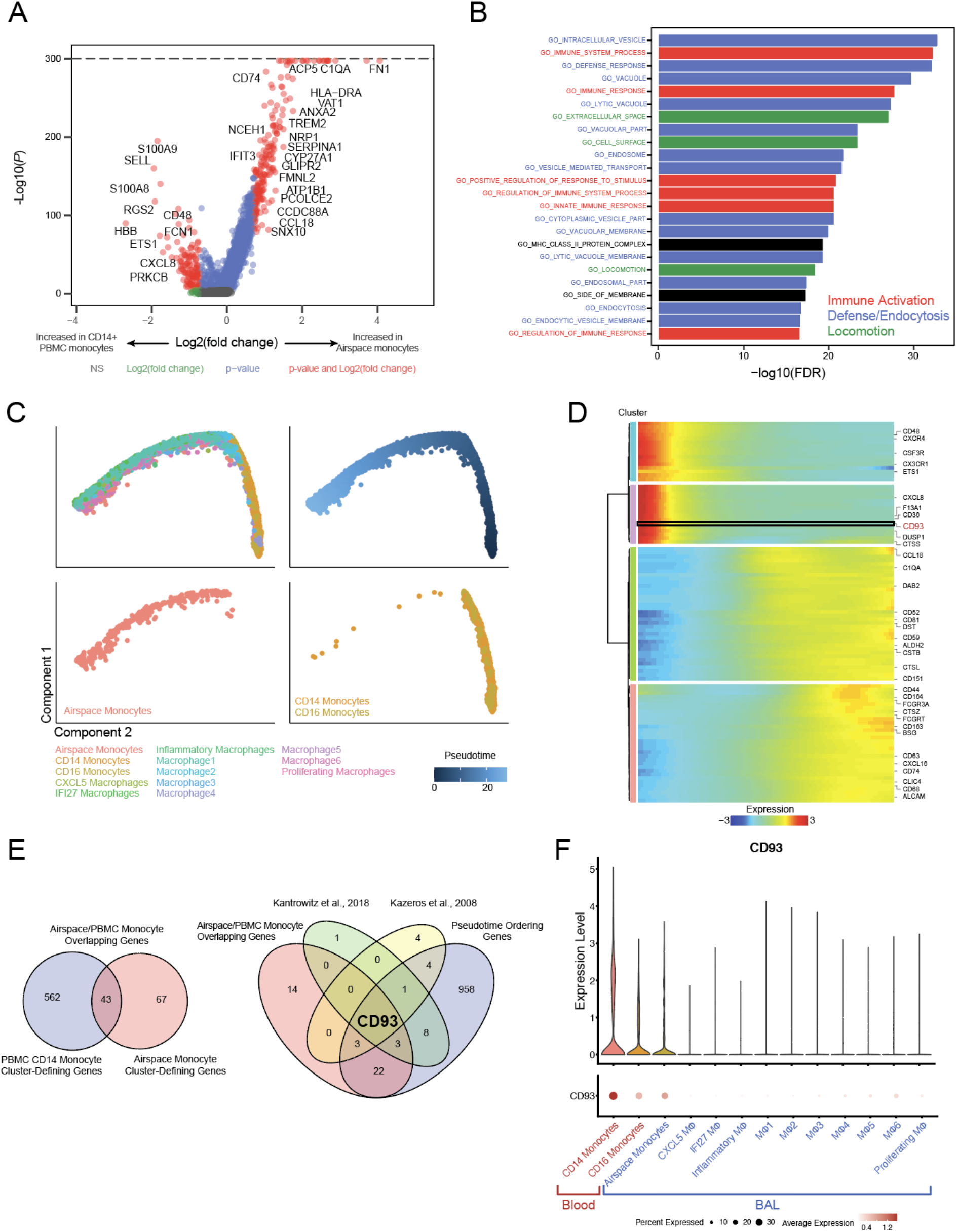
Airspace monocytes are distinct from blood monocytes but share the unique identifying marker CD93. (A) Volcano plot of differentially expressed genes between airspace monocytes and CD14+ PBMC Monocytes. X-axis represents airspace-vs-CD14+ monocyte fold change in expression (positive = increased in airspace monocytes; negative = increased in CD14+ PBMC monocytes), and y-axis represents Benjamini-Hochberg-corrected p-values. (B) Bar plot of gene programs enriched in airspace monocytes identified from GO enrichment analysis. (C) Pseudotime trajectory analysis of myeloid-lineage cell types from both blood and lung, colored by cell type (top left, bottom left, bottom right) or pseudotime coordinate (top right). (D) Heatmap of top genes differentially expressed along the pseudotime trajectory. (E) (Left) Venn diagram of the number and overlap of genes between airspace monocyte and CD14+ PBMC monocyte cell type-defining genes. (Right) Venn diagram of the number and overlap of genes analyses in four data sets. (F) Violin plot and dot plot of *CD93* expression across myeloid clusters.

To further investigate links between blood and airspace monocytes, we focused on the marker genes common to airspace and blood monocytes, thereby producing a gene signature which distinguishes monocytes in each compartment from mature AMs and MDMs. When we intersected airspace monocyte marker genes, blood monocyte marker genes, genes used to order the pseudotime trajectory, and marker genes from two bulk RNA-seq datasets of smoking-induced airway changes (*24, 25*), we found that *CD93* was the only gene which was shared independent of compartment, smoking status, analysis type, or data set (Figure 2E). Indeed, in our cohort, we found that *CD93* was expressed in blood and airspace monocytes but only sparsely detected by scRNA-seq in other myeloid cell types (Figure 2F). Concordantly, *CD93* is among the genes that are highest in blood monocytes and decrease with progression along the pseudotime trajectory towards mature macrophage states (Figure 2D). These data support airspace monocytes as a population derived from blood monocytes, infiltrating the airspaces, expressing CD93, and exhibiting an intermediate phenotype consistent with differentiation toward MDMs.

We next confirmed CD93 as a marker capable of identifying immature newly recruited monocytes in the airspaces at the protein level using flow cytometry. CD93 was not expressed by AMs but was expressed by 50% (median) of airspace monocytes, while 100% (median) of blood monocytes were positive for CD93 independent of smoking status (Figure 3A, Figure S5A). Given that CD14 and CD16 mark canonical blood and alveolar myeloid populations (i.e., CD14 as highly expressed in conventional blood monocytes, but decreased in mature alveolar macrophages; CD16 as only expressed on a subset of unconventional CD14^+/low^ blood monocytes but highly expressed by mature alveolar macrophages), we sought to evaluate how CD93 associated with these subtype- and maturation-linked markers (*26–28*). In contrast to CD93, CD16 was expressed by a small fraction of blood monocytes (median 9%), with a significantly higher proportion of airspace monocytes (median 75%) and all AMs (median 95%) (Figure 3B). CD93+ airspace monocytes expressed higher levels of *CD14* mRNA (associated with circulating blood monocytes) but lower levels of CD16 protein (otherwise associated with mature alveolar macrophages) than did CD93-airspace monocytes (Figure S5B-C). Thus, our data indicated that the airspace monocyte-identifying marker CD93 was downregulated or shed early during maturation while other markers of mature alveolar macrophages such as CD16 and CD14 were up- or down-regulated, respectively. To more directly explore marker levels during maturation, we isolated blood monocytes and followed CD93, CD16, and CD14 over a maturation period in human serum of nine days *in vitro*. Monocytes increased in size and granularity (SSC versus FSC) indicating their maturation state at different time points (Figure 3C). CD93 expression decreased rapidly 2h after adherence of blood monocytes and was almost undetectable after 24h of maturation (Figure 3D), while CD16 was significantly upregulated during the same time (Figure 3E); CD14 remained approximately constant (Figure 3F). We further sought to leverage our scRNA-seq atlases to demonstrate that CD93 enables additional specificity compared to the canonical myeloid markers of CD14 and CD16. In contrast to the specificity of CD93 for marking airspace monocytes among cells isolated through bronchoalveolar lavage and linking them to blood monocytes (Figure 2F), a variety of macrophage subsets expressed *CD14* only slightly lower than did airspace monocytes (Figure S5D), and airspace monocytes’ expression of *FCGR3A* (encoding CD16) was not substantially different from levels in mature macrophage subsets (Figure S5E). Thus, CD93 identifies an enriched monocyte population in the airspace of smokers, beyond the canonical myeloid markers of CD14 and CD16.

**Fig. 3.**
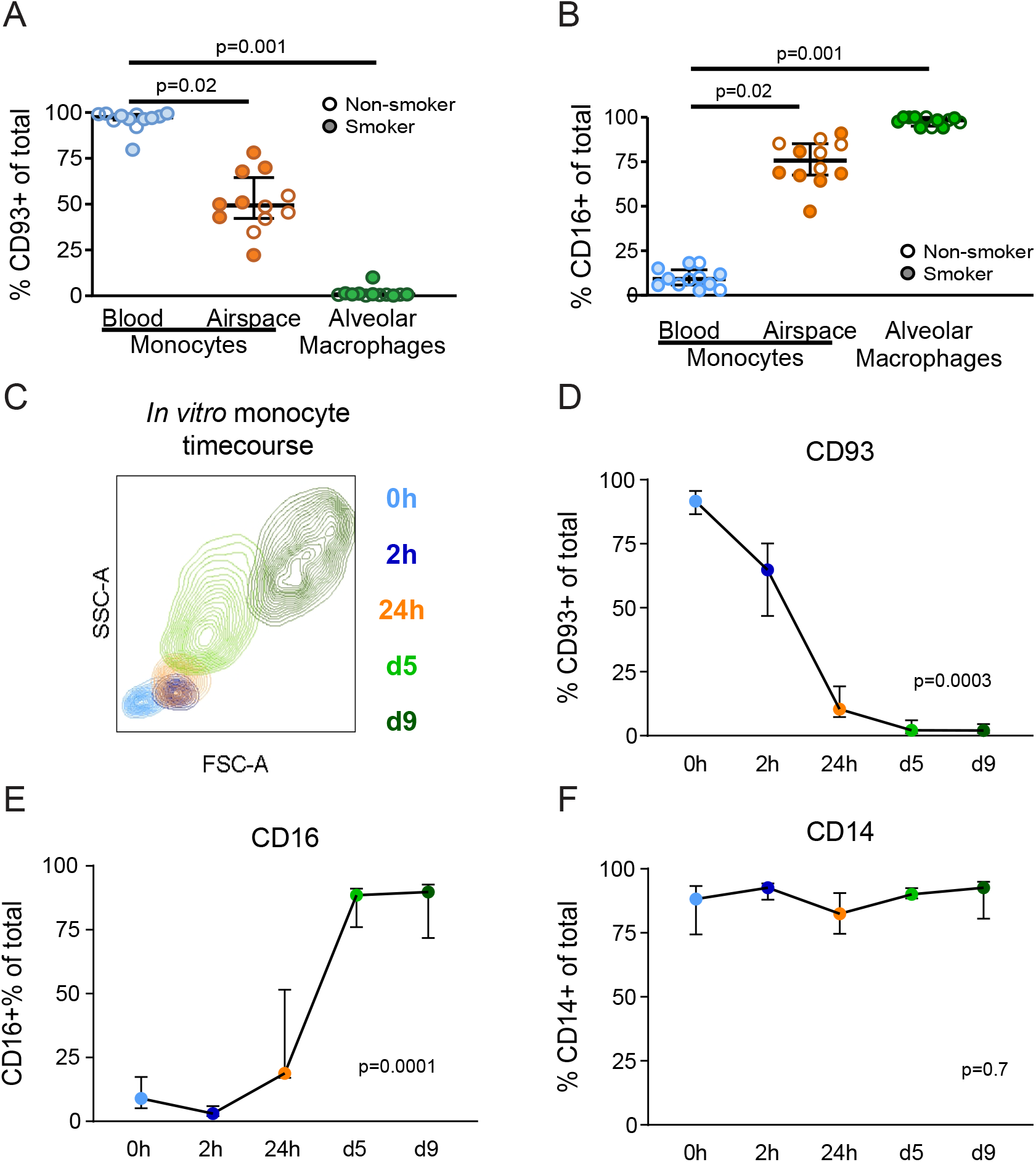
CD93 and CD16 reflect different stages of monocyte maturation *in vivo* and *in vitro*. Blood monocytes, airspace monocytes and macrophages were identified as shown in Figure 1 and analyzed for the expression of (A) CD93 and (B) CD16 using flow cytometry (smokers n=6; non-smokers n=5). (C-F) Blood monocytes were isolated from non-smokers (n=5) from PBMCs and differentiated into MDMs *in vitro*. MDMs were analyzed for (C) forward and side scatter distributions, (D) CD93, (E) CD16 and (F) CD14 expression at different time points using flow cytometry. The p-value was measured by (A and B) Kruskal-Wallis and Dunn’s multiple comparisons tests, or (D-F) Friedman test showing median with interquartile ranges.

In summary, airspace monocytes exhibit phenotypes intermediate between immature blood monocytes and mature macrophages. We identified CD93 as a unique marker for blood and airspace monocytes that is rapidly lost as airspace monocytes mature into macrophages and express markers including CD16, but provides additional specificity for delineating airspace monocytes from more mature macrophages in the alveolar environment. Therefore, CD14^+^CD93^+^CD16^-/low^ airspace monocytes represent a cluster of undifferentiated, recently transmigrated blood-derived monocytes and enriched in the lungs of tobacco smokers.

### Monocytes are recruited to the airspaces via CCR2-binding chemokines

Our analysis of myeloid cells revealed that tobacco smoking induces an accumulation of CD93+ airspace monocytes. Circulating monocytes express high levels of the chemokine receptor CCR2 and upregulate CCR5 upon transmigration and activation (*29*). To investigate the chemokines which recruit circulating monocytes into the airspaces, we measured 15 chemokines along with 10 cytokines in BAL fluid from smokers and never-smokers. We found significant correlations between the number of airspace monocytes and CCR2- and CCR3-binding chemokines (CCL2 and CCL7 as binding to CCR2; CCL11 as binding to CCR3; Figure 4A and Figures S6A-F). Although *CCR3* was only sparsely detected in scRNA-seq data, *CCR2* and *CCR5* were both significantly upregulated in airspace monocytes from smokers as compared to never-smokers, but not in blood monocytes (Figures 4B-C, S6G). We found recombinant CCL11 sufficient to attract monocytes in a CCR2-dependent manner *in vitro* (Figure 4D and Figure S6H); this CCR2-dependent transmigration was specific to monocytes, but was not observed for CD4+ or CD8+ T cells. Additionally, when treating BAL fluid with a blocking antibody against CCL11, transmigration of blood monocytes towards BAL fluid from smokers was significantly reduced, while a CCL2-blocking antibody did not produce a significant effect (Figure 4E). Thus, the recruitment of monocytes to the airspaces appears to involve smoking-associated increases in CCL11 in the lung.

**Fig. 4.**
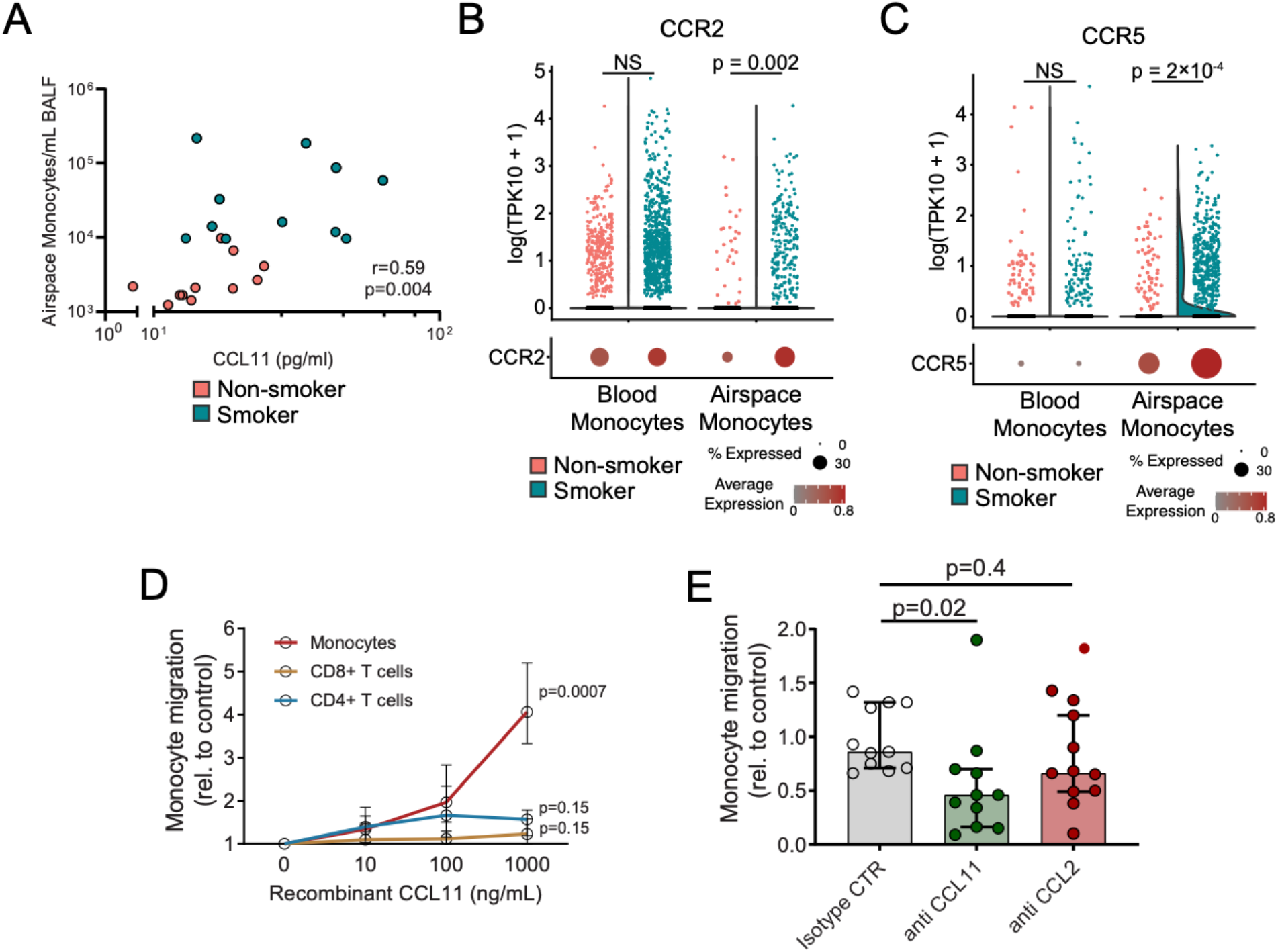
Non-classical chemokines recruit monocytes into the BAL of tobacco smokers. (A) BAL fluid (BALF) from smokers and non-smokers was concentrated (30x) and 15 chemokines measured using a multiplex system. CCL11 level in BAL fluid of smokers (turquoise circles) and non-smokers (salmon circles) correlated with the number of airspace monocytes / ml BALF using a Spearman non-parametric test. (B-C) Violin plot and dot plot of expression levels of the chemokine receptors (B*) CCR2* and (C) *CCR5* on airspace or blood monocytes from smokers (turquoise) or non-smokers (salmon). (D and E) PBMCs were layered on top of a 3μm transwell to monitor monocyte migration towards media containing (D) different concentrations of recombinant CCL11, or (E) concentrated BALF from smokers incubated with an isotype control antibody, antibodies against CCL11, or antibodies against CCL2. Statistical significance was evaluated by (A) Spearman’s correlation, (B-C) through a Wilcoxon rank-sum test with Benjamini-Hochberg correction for multiple hypothesis testing, or (D-E) Kruskal-Wallis and Dunn’s multiple comparisons tests showing median with interquartile ranges.

### Airspace and blood monocytes respond to *Mtb* with a strong inflammatory response compared to AMs

Monocytes can drive acute and chronic tissue inflammation, with an elevated inflammatory capacity as compared to that of macrophages (*30*). In a mouse model of human TB, uncontrolled infiltration of monocytes into the lung during infection was associated with increased pathology and susceptibility to accelerated *Mtb* growth (*31*). Further, the release of pro-inflammatory IL-1β is facilitated by an alternative pathway of inflammasome activation that has been shown to be active in monocytes but not in differentiated macrophages (*32*). Thus, we hypothesized that in the context of *Mtb* exposure, smoking-mediated recruitment of monocytes to the airspaces may result in an increased inflammatory milieu and a host lung microenvironment amenable to bacterial seeding and persistence. Compared to AMs, airspace monocytes were significantly enriched for pathways associated with immune activation, chemokine production, and chemotaxis, suggesting they have high inflammatory potential and may contribute to an accelerated inflammatory response in the lungs during infection (Figure 5A). Therefore, we tested whether monocytes and macrophages would respond with differential inflammatory responses *in vitro* when exposed to *Mtb*. Blood monocytes released a significant amount of IL-1β and other pro-inflammatory chemokines and cytokines after exposure to *Mtb*, while monocyte-derived macrophages (MDMs) and donor-matched AMs did not respond with a strong inflammatory response (Figure 5B and C; Table S3 and S4). However, adherent BAL cells from smokers, which contain ∼10-fold higher counts of airspace monocytes as compared to BAL samples from never-smokers (Figure 1B), secreted a significant amount of IL-1β and other pro-inflammatory chemokines and cytokines after exposure to *Mtb* (Figures S6I-J; note the scaling on y-axes across S6I-J; Table S3). In contrast, BAL cells from never-smokers largely did not produce a pro-inflammatory response following *Mtb* exposure. These results provide functional support of the pro-inflammatory transcriptional signatures observed in airspace monocytes and highlight how smoking may prime the lung immune environment for exaggerated inflammatory responses following pathogen challenge. To further characterize the inflammatory cells in BAL of smokers, we sorted airspace monocytes and AMs and compared the inflammatory response against donor-matched purified blood monocytes. Indeed, while AMs produced undetectable or low levels of inflammatory cytokines after exposure to *Mtb*, both matched airspace monocytes and blood monocytes secreted pro-inflammatory IL-1β upon *Mtb* stimulation, with airspace monocytes exhibiting intermediate levels of secretion between those of macrophages and blood monocytes (Figures 5D and E; Table S4). These results indicate that airspace monocytes have a heightened inflammatory phenotype after transmigration from blood into tissue, and therefore may represent the drivers of an accelerated inflammatory response following lung exposure to *Mtb*.

**Fig. 5.**
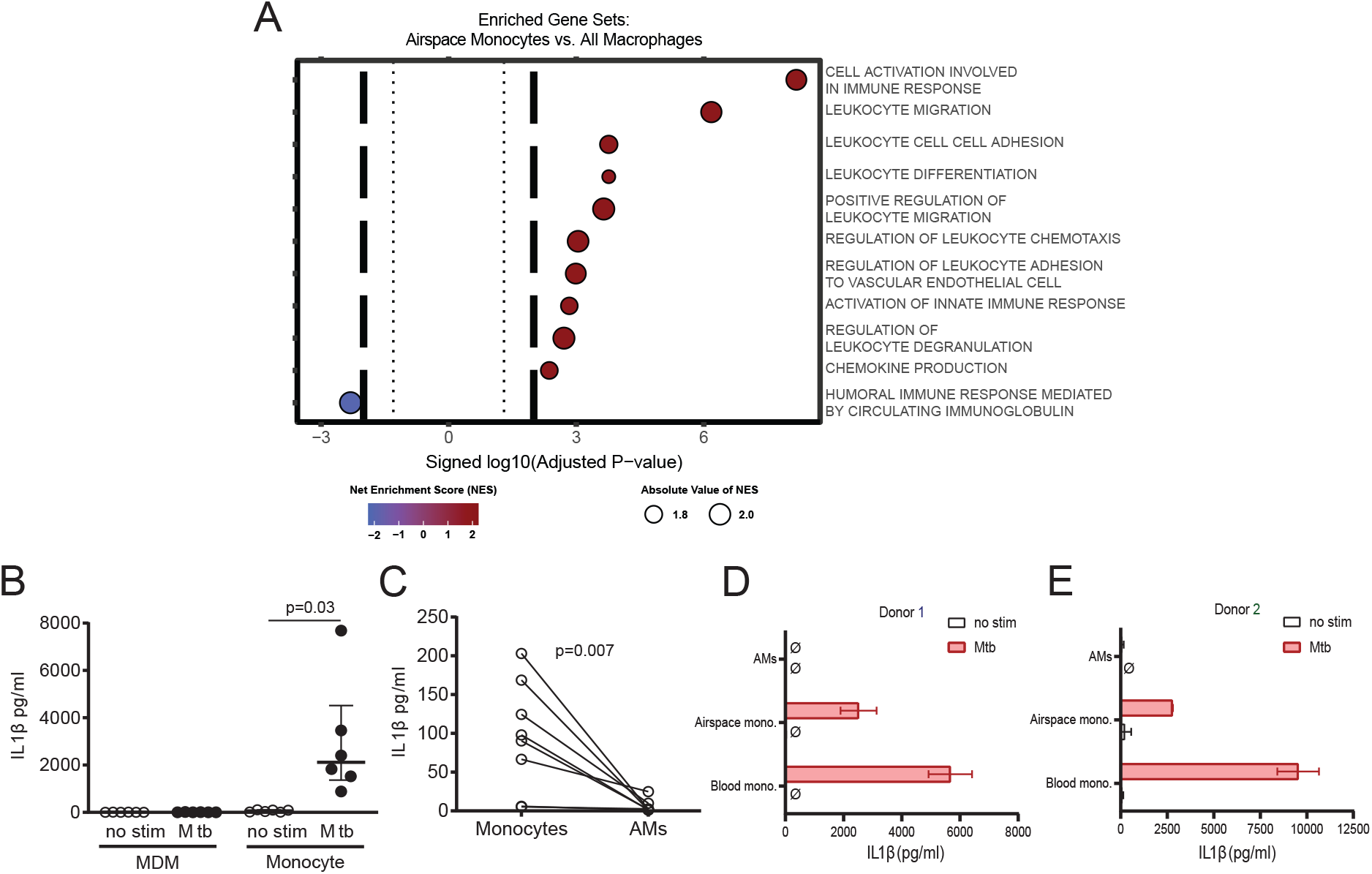
Blood and airspace monocyte are highly pro-inflammatory compared to mature macrophages. (A) GO processes enriched in GSEA analysis of differentially expressed genes between airspace monocytes and all macrophages. Benjamini-Hochberg correction was applied to p-values from GSEA analysis to control for multiple hypothesis testing. (B) IL1-β levels with or without 24 hours of *Mtb* exposure in monocyte-derived macrophages and blood monocytes. (C) IL1-β levels in donor-matched blood monocytes vs. BAL macrophages from non-smokers upon stimulation with *Mtb*. (D-E) IL1-β levels from donor-matched sorted blood monocytes, airspace monocytes, and BAL macrophages across 2 smokers. (B-E) Each dot represents a sample from different subjects. The p-values were measured by (B) Friedman test and Dunn’s multiple comparisons tests showing median with interquartile ranges. (D and E) bar graphs show mean of 2 technical replicates with standard deviation. Ø indicates that IL1-β was not detected.

### Monocyte inflammatory responses drive their susceptibility to *Mtb* growth

The accumulation of monocytes at the site of infection has been associated with increased susceptibility to *Mtb* growth in murine and zebrafish models (*33*). As such, the specific contribution of smoking to recruitment and enrichment of human airspace monocytes may represent a novel mechanism of accelerated growth of *Mtb*, thereby driving active TB in humans. However, direct links between tobacco-associated increases in airspace monocytes and *Mtb* intracellular growth have not been established. We found that *Mtb* growth was significantly increased in monocytes compared to mature MDMs (Figure 6A) or AMs (Figure 6B), but this intracellular growth was not associated with increased *Mtb* induced cell death (Figure S7A) *in vitro*. Adherent BAL cells from smokers, which contain a higher frequency of monocytes, showed a trend towards higher permissiveness to *Mtb* intracellular growth (Figure S7B), suggesting that BAL and blood monocytes are highly susceptible to intracellular *Mtb* growth. We therefore investigated if inflammatory monocyte states are specifically associated with increased susceptibility to *Mtb* growth. Monocytes and MDMs were treated with dexamethasone and the anti-inflammatory small molecule inhibitors SP600125 (Jun N-terminal kinase inhibitor). As expected, both inhibitors significantly reduced inflammatory cytokine production by monocytes following *Mtb* exposure (Figures S7C and D); however, they also inhibited *Mtb* intracellular growth specifically in monocytes (Figures 6C-F), supporting the role of pro-inflammatory monocytes in promoting *Mtb* growth.

**Fig. 6.**
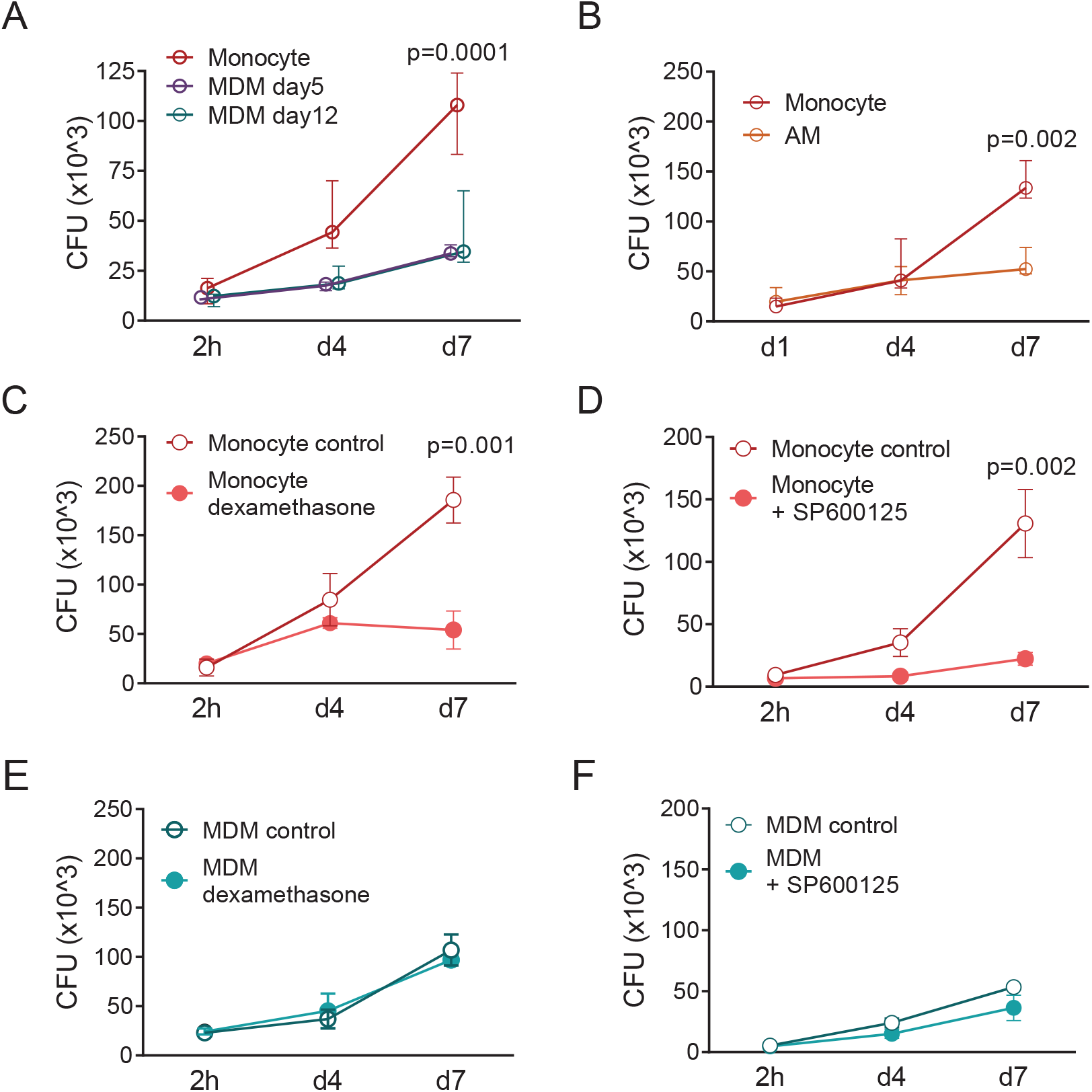
Elevated monocyte inflammatory response is associated with increased intracellular growth of *Mtb*. (A) Monocytes were isolated from human PBMCs (n=3) and infected with *Mtb* H37Rv at a MOI of 0.1 or differentiated for 5 or 12 days (MDM) prior to infection with *Mtb* H37Rv. At indicated time points macrophages were lysed to release intracellular bacteria and the number of intracellular bacteria was determined by counting colony forming units (CFU). MDM samples were slightly offset in the x-direction to aid with visualization. (B) AMs from non-smokers or donor-matched monocytes (n=4) were infected with *Mtb* H37Rv as described in A. (C-F) Monocytes or MDM were infected as described in A. Where indicated dexamethasone or SP600125 was added directly after 2h for the remaining time course. The p-values were measured by Friedman test and Dunn’s multiple comparisons test for each time point showing median with interquartile ranges.

Building on our observation that tobacco smoking drives increased recruitment of inflammatory immature monocytes to the airspaces, we found that the pro-inflammatory phenotype of these monocytes enables accelerated intracellular growth of *Mtb*, which can be abrogated through anti-inflammatory small molecule treatment *in vitro*. Thus, inflammatory immature airspace monocytes may represent a *Mtb*-permissive subpopulation that underlies the increased risk for active TB associated with smoking.

## DISCUSSION

Smoking has long been linked as a strong risk factor for active TB, yet the specific mechanisms underlying this observation remain incompletely understood (*5*). In this work, we performed a high-resolution analysis of lung immune cell composition, dynamics, and phenotypes in smokers to understand how tobacco smoking perturbs the lung microenvironment to promote increased *Mtb* growth. In our study, smoking was associated with enrichment of immature inflammatory airspace monocytes which likely were derived from newly recruited blood monocytes. We identified CD93 as a novel marker capable of distinguishing these immature airspace monocytes from more mature CD16^+^ airspace monocytes, AMs or MDMs, and provide evidence that CCL11, among other CCR2-binding chemokines, was an important chemoattractant involved in the specific recruitment of monocytes to the airspaces of smokers. Prior to *Mtb* exposure, airspace monocytes from smokers exhibited higher expression of inflammation- and activation-linked gene modules than did AMs. Consistent with this, BAL cells from smokers produced significantly elevated inflammatory responses after *Mtb* exposure as compared to cells from never-smokers, supporting the conclusion that smoking primes pro-inflammatory lung environments and immune responses. This increased inflammatory state was associated with enhanced *Mtb* growth, which could be inhibited by treatment with anti-inflammatory drugs (e.g., dexamethasone) specifically in monocytes, but not in macrophages, further supporting ties between smoking-induced immature inflammatory monocyte populations/states and susceptibility to *Mtb*. Overall, our data demonstrate that the increased recruitment and presence of immature inflammatory airspace monocytes may contribute to the increased risk of smokers to active TB.

We found an increased number of total BAL cells in smokers which was primarily explained by the infiltration of CD14+ monocytes. An increased number of myeloid cells in the airspaces of smokers has been reported before (*34–36*). However, in these studies, myeloid cells were identified using methods that did not discriminate between monocyte and macrophage subpopulations. One study found a decrease of CD16 expression and slight increase of CD14 expression on the surface of BAL cells from smokers and COPD patients, consistent with the CD14+ monocyte phenotype in our cohort (*35*). Intriguingly, using scRNA-seq, characterization of airspace monocytes which differentiate further into a number of macrophage phenotypes has also been described in patients with COVID-19 and COPD, confirming the connection between inflammation, disease, and recruited monocytes across multiple disease contexts (*18, 37–39*). We specifically identify CD93 as a novel marker for defining a disease-relevant population of airspace monocytes, enriched in smokers’ airspaces. Importantly, we also demonstrate that CD93 enables more specific identification of these pro-inflammatory airspace monocytes as compared to canonical myeloid markers. Thus, CD14^+^CD93^+^CD16^low^ monocytes in the airspaces represent an intermediate state between the transcriptional and functional characteristics of blood monocytes and AMs, with airspace monocytes importantly retaining pro-inflammatory phenotypic characteristics of blood monocytes despite translocation into the tissue microenvironment. Examination of myeloid-associated disease-linked gene sets, RNA velocity, and pseudotime analyses all supported the observation that airspace monocytes differentiate towards more mature macrophage states. Furthermore, *ex vivo Mtb* exposure assays demonstrated that the protein-level functional responses of airspace monocytes are capable of driving inflammation in the airspaces. Thus, our work demonstrates smoking-mediated recruitment of immature inflammatory monocytes that retain functional characteristics of blood monocytes despite ongoing differentiation, thereby establishing a lung niche conducive to elevated *Mtb* growth.

As immune intercellular signaling patterns underlying airspace monocyte recruitment, our data highlights CCL11-CCR2 signaling as key for driving monocyte migration (with additional chemokines also being correlated with airspace monocyte counts). CCL11 is best known in the context of allergic asthma and eosinophil recruitment via CCR3 (*40*). However, it can also bind CCR2 (*41*) which is highly expressed on blood monocytes (*42*) and may facilitate myeloid cell recruitment, which had previously been shown in atopy (*43*), suggesting pleiotropic effects important in smoking-induced, airspace monocyte-linked *Mtb* susceptibility. However, in asthma, CCL11 requires a Th2 milieu and IL5 for effective eosinophil recruitment (*44*). In our cohort of smokers, we did not detect a bias towards type 2 immune response and consequently did not observe any significant increase in infiltration of eosinophils or neutrophils in airway samples. Instead, we detected an accumulation of pro-inflammatory airspace monocytes as associated with CCL11-CCR2 signaling. Building on these findings, future studies with larger cohorts could evaluate significant shifts in chemokine concentrations in the air spaces and plasma in never-smokers versus smokers.

Furthermore, we found a link between the pro-inflammatory phenotypes of airspace monocytes and increased *Mtb* growth, highlighting the importance of regulated, balanced immune responses to pathogen control. Innate immune activation and mobilization of a hybrid type 1/type 17 adaptive immune responses are crucial for successful *Mtb* control (*45*); however, aberrant pro-inflammatory states can also be detrimental to the host (*46, 47*). Prior work has suggested that elevated infiltration of immature inflammatory monocytes into the lung is associated with inflammation-induced tissue damage and higher susceptibility to *Mtb* (*31, 48, 49*). This model is further supported by findings demonstrating that highly pathogenic *Mtb* strains can drive increased pathology via elevated infiltration of monocytes (*33*). Protective roles of pro-inflammatory immune responses in *Mtb* may depend on timing, location, and concentration of released signaling molecules, such that stronger inflammatory responses may not necessarily result in improved *Mtb* control. In fact, we found that treatment of monocytes with the anti-inflammatory molecules dexamethasone or SP600125 resulted in reduced *Mtb* intracellular growth. Interestingly, the two anti-inflammatory mediators used in this study have relatively broad mechanisms of action. Dexamethasone inhibits inflammasome activation, while SP600125 acts as a Jun N-terminal kinase inhibitor. Given the effectiveness of both drugs in inhibiting *Mtb* growth in our *ex vivo* functional assays, their effects on *Mtb* growth would not seem to be inflammasome-dependent. We also note that dexamethasone and similar adjunctive glucocorticoids are already approved to reduce inflammation and increase survival in patients with tuberculosis-associated immune reconstitution inflammatory syndrome (TB-IRIS) and TB associated meningitis (*50, 51*). Our data suggest there may also be a benefit during treatment of inflammatory diseases in tobacco smokers or patients with similarly pro-inflammatory lung environments.

A limitation of this study is relatively small sample sizes. Future work could seek to replicate these results in larger cohorts. Towards this end, we were able to support our findings through comparisons against other previously published studies on lung diseases and smoking. Such analyses revealed not only concordant phenotypes across diseases and cohorts, but also that our findings are not driven by confounders related to cohort characteristics. Further meta-analyses leveraging published scRNA-seq datasets and mouse models of human disease (*52*) could enable broader inquiries into mechanisms of smoking as a risk factor for other pulmonary diseases via identification of similar recruited human airspace monocyte populations. Likewise, while we demonstrated that smoking leads to increased recruitment of immature inflammatory monocytes to the airspaces and primes a niche for *Mtb* growth, further studies are needed to define more precise mechanisms of how these pro-inflammatory monocytes permit rapid *Mtb* growth. Smoking has been reported to impair the ability of AMs to control *Mtb* intracellular growth (*6, 9, 11*). Our data suggest that recruitment of immature inflammatory monocytes into the airspaces additionally provides a highly permissive cell type for *Mtb* growth that is also capable of inducing broader lung inflammation. Thus, our findings indicate that airspace monocytes provide an additional independent mechanism by which tobacco smoking increases the risk of *Mtb* acquisition and progression to active TB. Epidemiological studies have not fully elucidated whether smoking-associated active TB cases and TB mortality are due to increased risk of primary infection, re-activation or re-infection. However, in locations with high TB prevalence and accompanying elevated likelihood of re-infection (e.g., as compared to re-activation), smoking-induced airspace monocytes might increase the risk of accelerated growth and dissemination of *Mtb* after repeated exposure.

Our work demonstrates how environmental exposures can drive dysregulation of tissue microenvironments that in turn impacts susceptibility to disease. Additionally, we identify potential translational directions and specific small molecule approaches to modulate immune function for improved TB outcomes. These findings provide new biological understanding of the mechanism by which smoking increases *Mtb* growth and reveal potential strategies to mitigate TB burden among tobacco smokers.

## MATERIALS AND METHODS

### Study design

Detailed inclusion and exclusion criteria and the study protocol are provided in the supplemental material and methods. Smokers and never smokers were defined according to CDC guidelines as current smokers (every day smokers; n=13) with ≥ 100 lifetime cigarettes and never-smokers (n=21) who never smoked or smoked less than 100 cigarettes in his or her lifetime (Table S1 and supplementary material and methods). Eligible subjects underwent bronchoscopy and phlebotomy. Pulmonary function testing (PFT) was performed for all participants, and were within normal limits (IQR of FEV1 and FVC ≥ 80%). The study was approved by the Partners Healthcare Institutional Review Board. Subjects gave their written informed voluntary consent prior to inclusion in the study.

### Processing of Human Samples

Broncho-alveolar lavage (BAL) fluid was obtained from the right middle lobe and lingula by washing each with 120 ml of saline delivered in 4 x 30 ml aliquots. Samples were combined, filtered through a 70μm cell strainer and centrifuged at 4°C for 10min at 400xg before resuspending cells in D-PBS (Gibco) determining cell counts and viability using a nucleocounter®. PBMCs were obtained from whole blood EDTA samples and by density centrifugation using histopaque® (Sigma). Samples were selected based on cell viability (at least 80% or higher) and cell numbers to perform all assays (at least 5×10^6^ total cells). Samples for scRNAseq (non-smoker n=4; smokers n=5) were selected based on sample availability and viability. All BAL and PBMCs were used freshly for downstream analysis and assays. The BAL fluid was aliquoted and frozen at -80°C until further usage.

### Flow cytometry

Cells from BAL or PBMCs were stained with fluorescent-conjugated antibodies and fixable viability dye. For this, cells were washed in FACS buffer (D-PBS (Gibco) + 1% FCS + 5 mM EDTA (Gibco)) and incubated with 5μl TruStain FcX receptor blocking solution (Biolegend) for 5min at RT, before staining with antibodies for 10min at RT followed by viability staining for 5min at RT. Cells were washed again with MACS buffer before fixation in 4% PFA. Details for all used antibodies are provided in Table S2. Samples were run on an LSR Fortessa (BD Biosciences, San Jose, CA, USA) and analyzed using FlowJo (BD Biosciences).

### Cytokine and chemokine detection

BAL fluid was thawed and 30-fold concentrated using ultra centrifugal filter units (Millipore). Chemokines and cytokines were measured with a customized Milliplex panel (EMD Millipore) on a Luminex-200 and analyzed using xPONENT 3.1 software (Luminex Corp.). Some of the analyzed data have been analyzed and presented in a previous publication (*12*).

Bulk BAL cells or FACS sorted airspace monocytes, AMs and blood monocytes from PBMCs from two smokers (Donor 1 and Donor 2) were used to determine the inflammatory response of myeloid cells to *Mtb*. 50,000 cells per cell type were plated into a 96 well plate (in duplicates) and either left untreated as a control or exposed to *Mtb* at an MOI of 1 for 24h.

The supernatants from each well were collected, sterile filtered and used in a Milliplex assay to measure cytokines and chemokines in each sample using a 5- and 25plex customized kit (Millipore and R&D Systems). Data were analyzed by calculating the average value of the biological duplicates and then the ratio of *Mtb* treated / control. This was done for each of the measured cytokines. The data were grouped for each sample type (airspace monocyte, AMs and Blood monocytes) and a Kruskal-Wallis with Dunn’s multiple comparisons test to test for significance between airspace monocytes vs AMs and Blood monocytes vs. AMs.

### Monocyte *in vitro* differentiation

Monocytes were purified from freshly isolated PBMCs by CD14 positive selection using MACS® cell separation (Miltenyi), washed with MACS buffer before counting and plated out at 0.1×10^6^ CD14+ cells per well in a 96 well plate and RPMI medium (Gibco) containing 10% human serum for differentiation into MDMs for 5 days unless otherwise indicated.

### Transwell assay

Monocyte *in vitro* migration towards BAL fluid or recombinant CCL11 (R&D Systems) was determined in a 3μm transwell chamber (Corning) by placing PBMCs into the upper well and counting CD14+ viable monocyte in the lower chamber after 3h by flow cytometry. Where indicated PBMCs were pre-treated with anti-CCR2 (8μg/ml, clone 48607, R&D Systems), anti-CCR3 (8μg/ml, clone 61828, R&D Systems) or anti-CCR5 (5μg/ml, clone 45531, R&D Systems) blocking antibodies, or BAL fluid was pre-incubated with an anti-CCL11 (10μg/ml, clone L403H11 Biolegend), anti-CCL2 (10μg/ml clone 2H5 Biolegend), or isotype control antibody (rat IgG1, 10μg/ml; Biolegend) 30min before the transwell assay.

### *Mtb* culture and *in vitro* infection

Viable *Mycobacterium tuberculosis* H37Rv were used from a freshly thawed stock after expansion during exponential growth phase in 7H9 medium (Middlebrook). The bacteria aliquot was centrifuged at 1200xg for 5 min and resuspended in PBS before photometric determination of bacterial numbers. The culture was diluted for a MOI of 0.1 in RPMI + FCS and added to 0.1×10^6^ monocytes or macrophages in a 96 well plate. After 2h extracellular bacteria were washed off three times with PBS before adding back RPMI + 10% FCS medium. Where indicated, cells were treated with dexamethasone (0.1µM) or SP600125 (0.1µM) after uptake of the bacteria and for the remaining time course. At indicated time points, intracellular bacteria were released by incubation of each well with 0.05% Triton-X (Sigma) in 200μl in PBS for 5min. Colony forming units (CFU) were determined by plating out the Triton-X treated supernatants on 7H11 agar plates (Middlebrook) in at least 2 serial dilutions and counting the CFU plates after 3-4 weeks incubation at 37°C in the dark.

### scRNA-seq Methods

scRNA-seq was conducted using Seq-Well S^3^ protocol (Supplemental Material and Methods). In brief, individual cells were loaded in nanowells with poly-dT capture beads. Following RNA capture, reverse transcription, second-strand synthesis, and PCR amplification, sequencing libraries were constructed using the Nextera XT DNA Library Preparation Kit. Samples were sequenced on a NovaSeq 6000 and aligned to the Hg19 genome on the Broad Institute’s Terra cloud computing platform. Cells were filtered based on common quality metrics, followed by downstream analyses in Seurat v.4.0.1. RNA velocity analyses were conducted in scVelo and Scanpy, and pseudotime analyses with Monocle 2. Gene set enrichment analyses were conducted using fgsea (version 1.16.0).

### Statistical Analysis

All statistical analysis except for scRNA-seq was performed using GraphPad Prism. For scRNA-seq, Seurat v.4.0.1, scVelo, Scanpy, and Monocle 2 were used for analyses, with Seurat v.4.0.1 and ggplot2 used for plotting. DirichletReg was used to implement Dirichlet regression for scRNA-seq compositional analyses. Data are expressed as median and interquartile range unless otherwise noted. Each figure legend contains details about the specific statistical test. Differences were considered statistically significant when P < 0.05 after correction for multiple hypothesis testing.

## Acknowledgments

The authors would like to acknowledge the contributions of the Ragon Institute Imaging Core Facility. The authors also thank the staff from the Massachusetts General Hospital. We would also like to thank F. Ruzicka, S. Ip, D. Worrall, M. Kone, V. Kelly and the Pulmonary Special Procedures Unit for their assistance with this study.

## Funding

National Institutes of Health grant NIH 1U01HL121827-01 (BDM and DSK)

National Institutes of Health grant NIH 1R01DA042685-01 (DSK)

National Institutes of Health grant NIH 5U24AI118672 (AKS)

National Institutes of Health grant NIH DP2OD020839 (AKS)

Ragon Institute of MGH, MIT, and Harvard (AKS)

Beckman Young Investigator Award (AKS)

Alfred P. Sloan Research Fellowship in Chemistry (AKS)

National Science Foundation Graduate Research Fellowship (1122374) (CNT)

Fannie and John Hertz Foundation Fellowship (CNT)

## Author contributions

Conceptualization: BC, JLC, BDM, AKS, DSK

Methodology: BC, CNT, MHW, JLC, BDM, AKS, DSK

Investigation: BC, CNT, MHW, JLC, AHL, AES, AKD

Visualization: BC, CNT, MHW

Funding acquisition: BDM, AKS, DSK

Project administration: JLC, AHL, AES, AD

Supervision: BC, BDM, AKS, DSK

Writing – original draft: BC, CNT

Writing – review & editing: BC, CNT, MHW, JLC, BDM, AKS, DSK

## Data and materials availability

All data associated with this study are present in the paper or the Supplementary Materials. All codes used for scRNAseq analysis are accessible under XX. All sequencing files are available at Gene Expression Omnibus (GEO) repository under XX.

## Supplementary Material

### Material and Methods

Detailed Inclusion and Exclusion Criteria

Volunteer subjects were recruited via approved advertisement.

### Inclusion Criteria

1. Male or female, between 18 to 65 years of age
2. Laboratory values within 45 days prior to enrollment that meet the following criteria:
a. Hemoglobin ≥ 10.0 g/dL
b. Absolute neutrophil count ≥ 1000/mm3
c. Platelet count ≥ 80,000/mm3
d. Prothrombin time (PT) < 1.5 x upper limit of normal (ULN)
e. Partial thromboplastin time (PTT) < 1.5 x ULN
3. Negative urine pregnancy test (sensitive to 25 IU HCG) at screening and within 24 hours of the study procedure for female participants who are able to get pregnant
4. Ability and willingness to give written informed consent and to comply with study requirements

### Exclusion Criteria

1. History of clinically significant pulmonary disease including but not limited to asthma, chronic obstructive pulmonary disease, pulmonary fibrosis, bronchiectasis, or pulmonary hypertension
2. Female subject who is pregnant or less than 8 weeks post-partum
3. Use of any immunomodulatory agents within 30 days prior to study enrollment
4. History of underlying medical condition for which antibiotic prophylaxis for invasive procedures is required
5. History of intolerance, sensitivity, allergy or anaphylaxis to benzodiazepines or other narcotics to be used during the procedure
6. History of intolerance, sensitivity, allergy or anaphylaxis to lidocaine or other amide anesthetics, as well as benzocaine or other ester type anesthetics
7. History of coronary artery disease, myocardial infarction, chronic renal failure, decompensated cirrhosis, or any other condition that in the opinion of the investigator will compromise ability to participate in the study
8. Currently taking anticoagulants including but not limited to: heparin (Hep-Lock, Hep-Pak, Hep-Pak CVC, Heparin Lock Flush), warfarin (Coumadin), tinzaparin (Innohep), enoxaparin (Lovenox), danaparoid (Orgaran), dalteparin (Fragmin), clopidogrel (Plavix), prophylactic aspirin, and regular NSAID use
9. Currently taking any of the following medications: systemic steroids, interleukins, systemic interferons, or systemic chemotherapy
10. Systemic antibiotic therapy within 30 days of enrollment or procedure
11. Currently employed at or affiliated with Ragon Institute of MGH, MIT, and Harvard

### Study Protocol

#### Screening Visit

The screening visit included review of the pre-screening questionnaire, a complete medical history and review medical records, a pulmonary function test (PFT), and a physical examination. Female subjects of childbearing potential underwent urine pregnancy testing. All subjects underwent phlebotomy and spirometry. Phlebotomy was performed to obtain T cells counts, safety labs (PT, PTT, complete blood count with differential) and a blood sample for research purposes. Subjects > 55 years of age also had an electrocardiogram.

#### Bronchoscopy Visit

At the bronchoscopy visit, female subjects of childbearing potential underwent repeat urine pregnancy testing. A blood sample for research purposes was also collected. Topical anesthesia was achieved using topical lidocaine (≤300 mg). Flexible bronchoscopy was then performed under conscious sedation. Bronchoalveolar lavage (BAL) fluid was obtained by washing 120 ml of saline (4 x 30 ml aliquots) sequentially in a segment of the lingula and the right middle lobe for a total volume of 240 ml. BAL fluid was collected into sterile containers and stored on ice until processing.

### LDH assay

Supernatants were collected from infected and uninfected myeloid cell cultures and frozen in aliquots at -80°C. Freshly thawed supernatants were used in the LDH assay following the manufacturer’s protocol (Roche). As a positive control serial dilutions of Triton-X 100 (Sigma) treated myeloid cells matched for cell type and cell numbers were used to calculate the frequency of dead cells.

### Single-Cell RNA-sequencing (scRNA-seq) Processing, Sequencing, and Alignment

Massively-parallel scRNA-seq was performed using the Seq-Well S^3^ platform as described in Hughes et al. In brief, a functionalized polydimethylsiloxane (PDMS) array was loaded with uniquely barcoded mRNA capture beads (ChemGenes; MACOSKO-2011-10), then 10-15,000 cells. After cells settled into wells, a hydroxylated polycarbonate membrane with 10 nm pore sizes was used to confine biological molecules within wells while allowing buffer exchanges for cell lysis and mRNA transcript hybridization to beads. Beads were removed from the array and pooled for reverse transcription, followed by Exonuclease I treatment on the obtained cDNA product to remove excess primer. Second-strand synthesis was performed to produce double-stranded cDNA, followed by PCR amplification. Sequencing libraries were prepared using the Nextera XT DNA Library Preparation Kit. Version 2 of the Drop-seq pipeline (previously described in Macosko et al.) was used for sequencing read alignment. Bcl2fastq was used to convert raw sequencing reads from bcl files to FASTQs, based on Nextera N700 indices for individual samples. STAR and the DropSeq pipeline were used to align demultiplexed FASTQs to the Hg38 genome on the Broad Institute’s Terra cloud computing platform. Read 1 of each sequencing fragment contained a 12-bp barcode and 8-bp unique molecular identifier that tagged each individual read. After alignment, the 12-bp cell barcodes were used to group reads, followed by collapsing reads based on 8-bp UMI to generate digital gene expression (DGE) matrices.

### scRNA-seq Data analysis

#### Cell Quality Filtering

Cells were first filtered based on the number of detected genes (>500 genes per cell). On a sample-by-sample basis, we calculated variable genes and conducted principal component analysis, retaining the top 30 components that were also identified as significant with jackstraw simulations. UMAP dimensionality reduction and clustering were performed using the significant components. Within each sample, clusters distinguished by high proportions of mitochondrial gene expression were removed from downstream analyses, as these correspond to low-quality cells. Clusters were identified both through marker discovery (i.e., likelihood-ratio test, Bonferroni-corrected) and manual curation with prior literature.

#### RNA Velocity

RNA velocity analyses were conducted using scVelo (version 0.2.3). Briefly, the neighborhood graph was calculated with scanpy.pp.neighbors, followed by moments for velocity estimation using scv.pp.moments. RNA velocities were then calculated using scv.tl.velocity in deterministic mode, followed by calculation and visualization of RNA velocity in UMAP space using scv.tl.velocity_graph and scv.pl.velocity_embedding_grid, respectively.

#### Pseudotime

The pseudotime analyses were performed using Monocle2. Briefly, cell size factors and gene dispersions were calculated with estimateSizeFactors and estimateDispersions, respectively. Genes were filtered to those above an expression threshold of 0.1 and detected in at least 10 cells; cells with outlier numbers of UMIs (defined as more than 2.5 standard deviations away from the mean UMI count across all cells, on a log10 scale) were likewise filtered. Gene expression values were log-transformed and standardized. Cells were ordered based on genes differentially expressed across cell types (determined via differentialGeneTest), via reduceDimension (method = “DDRTree”) and orderCells.

#### Compositional Analyses

For determining compositional changes with smoking status while accounting for compositional shifts in one cell type affecting relative proportions of other cell types, we aggregated cell type counts by patient and implemented Dirichlet regression using the DirichletReg package (version 0.7-1).

#### Gene Set Enrichment Analyses

For determining enrichment of gene sets across cell states and biological conditions, genes were ranked by log-fold change between the two conditions of interest, followed by gene set enrichment using fgsea (version 1.16.0). msigdbr (version 7.2.1) was used to access GO processes (category C5, subcategory GO:BP).

#### Gene Set Module Scoring

AddModuleScore (as implemented in Seurat, version 4.0.1) was used to determine cell clusters’ expression levels of gene sets, for downstream comparisons and correlations. To further define Airspace monocyte phenotypes in this study, marker gene lists for the “monocyte-like macrophage” cell state and “lipid metabolism-associated genes” were used from Baβler et al. To evaluate the effects of age and biological sex covariates on airspace monocyte phenotypes, we used lists of genes upregulated in alveolar macrophages with age from Angelidis et al.; genes upregulated with age in lung cells from Chow et al.; genes upregulated in alveolar macrophages with smoking from Morrow et al.; and genes on autosomes (i.e., not on the X or Y chromosome) associated with male or female biological sex in lung cells from Yang et al. (subset to only include genes on autosomes).

## Supplementary Figures

**Fig. S1.**
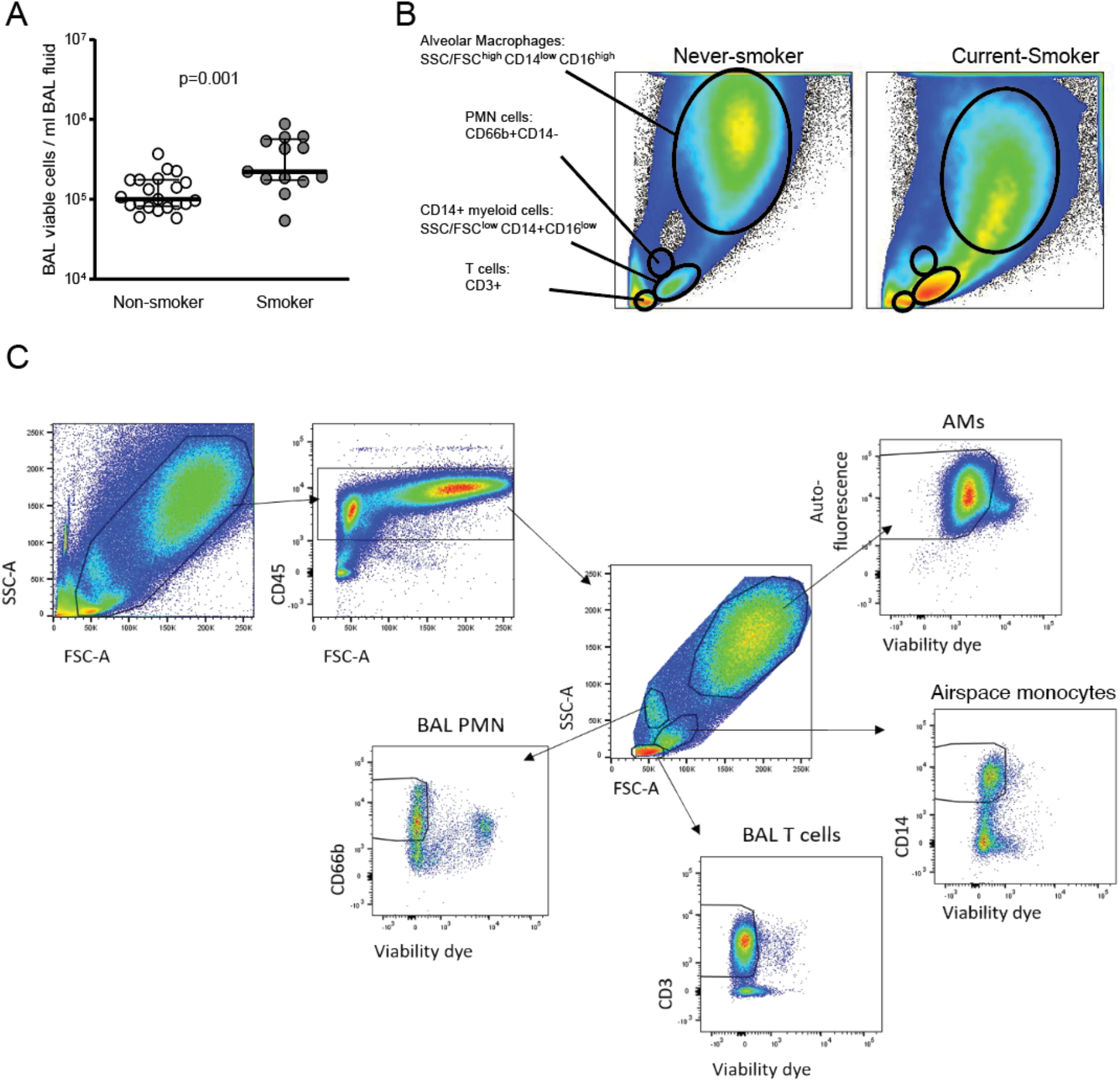
Analysis of BAL cells by flow cytometry. (**A**) The total number of viable BAL cells from non-smokers and smokers was assessed using a NucleoCounter. (**B and C**) Gating strategy to identify cell populations in BAL by flow cytometry. CD45+ cells were further analyzed as AMs (SSC/FSC^high^ autofluorescence^high^ fixable viabilty dye^-^, airspace monocytes (SSC/FSC^low^ CD14^+^ fixable viability dye^-^), BAL T cells (SSC/FSC^low^ CD3^+^ fixable viability dye^-^), and BAL PMN (SSC/FSC^low^ CD66b^+^ fixable viability dye^-^).

**Fig. S2.**
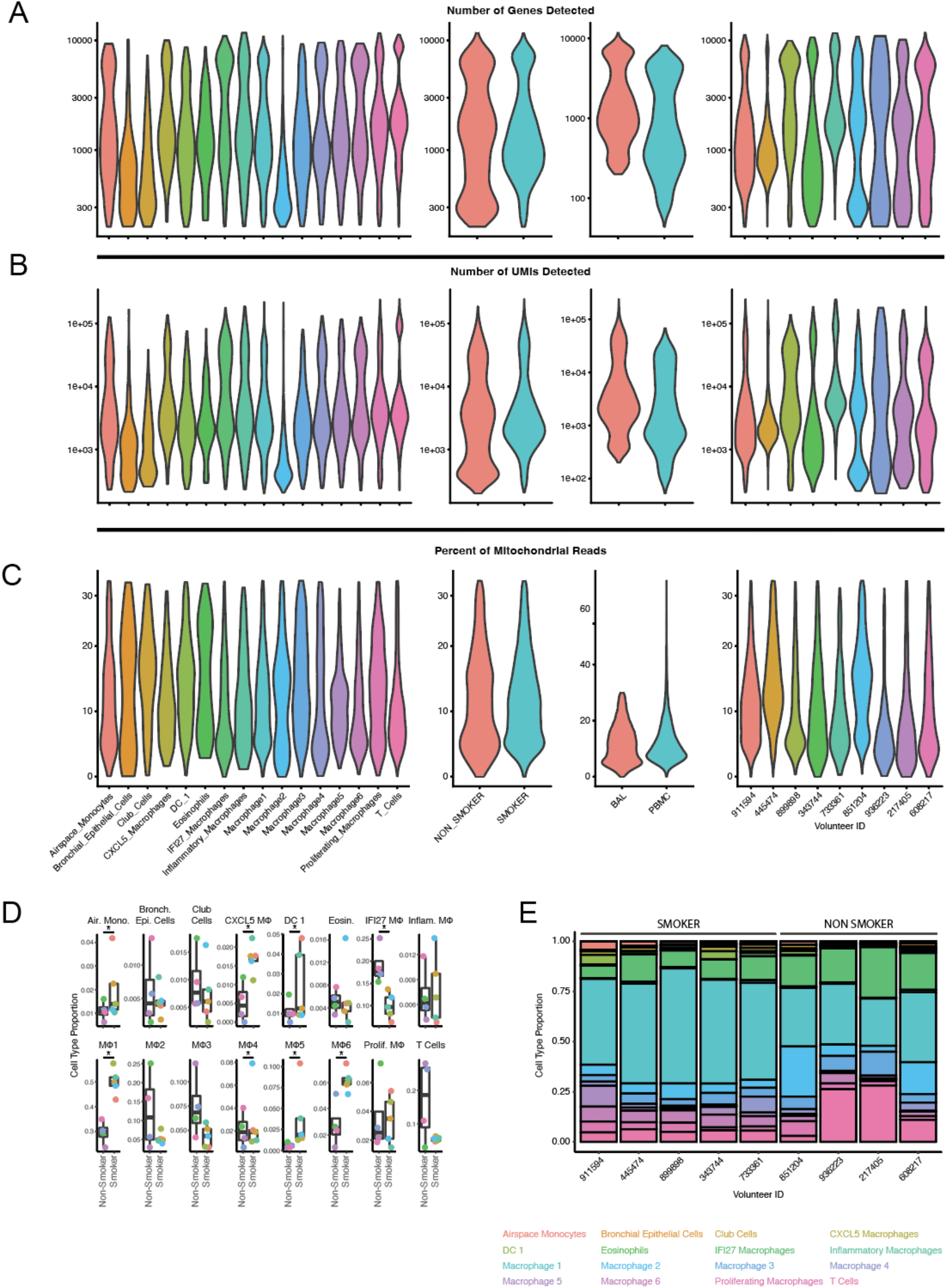
Quality metrics of scRNA-seq dataset. **(A)** Distribution of genes detected per cell, split by cluster identity (far left), smoking status (middle left), compartment (middle right), and volunteer (far right). **(B)** Distribution of unique RNA molecules detected per cell, split by cluster identity (far left), smoking status (middle left), compartment (middle right), and volunteer (far right). **(C)** Distribution of percent mitochondrial RNA molecules detected per cell, split by cluster identity (far left), smoking status (middle left), compartment (middle right), and volunteer (far right) **(D and E)** Composition of each cell type in smoking vs. non-smoking volunteers (Dirichlet regression analysis, which accounts for compositional shifts in one cell type necessarily affecting relative proportions of other cell types; * indicates p < 0.05).

**Fig. S3.**
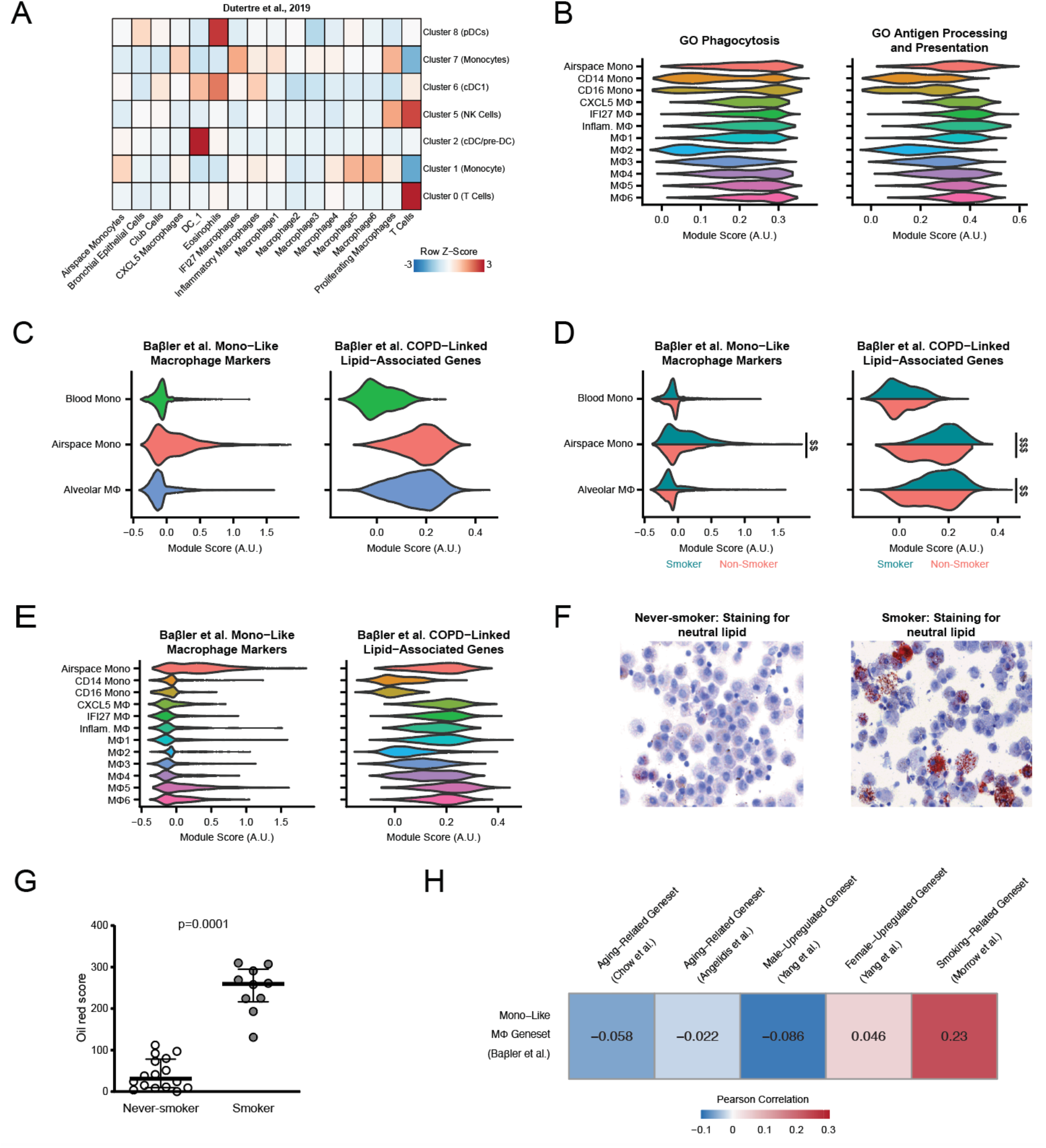
Links Between Transcriptional Programs, Cellular Identities, and Volunteer Demographic Characteristics. **(A)** Heatmap of scoring clusters from this study against markers derived from previously published blood immune atlas from Dutertre et al. **(B)** Module scoring for GO processes relevant to monocyte/macrophage function, split by cluster identity. (**C**) Module scoring for gene lists from Baβler et al., split by broad myeloid division. (**D-E**) Module scoring for GO processes and gene lists from Baβler et al. (Frontiers in Immunology, 2022), showing differences based on smoking status within broad myeloid divisions. Effect size measured as Cohen’s D: $ 0.2 < D < 0.5; $$ 0.5 < D < 1; $$$ D > 1. (**F-G**) BAL cells from non-smokers and smokers were placed onto a glass slide using a cytospin centrifuge and stained with Oil red to identify lipid loaded foamy macrophages. The p-value was measured by a Mann-Whitney test showing median with interquartile ranges. (**H**) Pearson Correlation between airspace monocyte-associated phenotype (module scoring for Baβler et al.’s Monocyte-Like Macrophage gene set) and gene sets of transcriptional differences associated with age, biological sex, and smoking in the lung. The cohort had more males than females (Table S1), so that if the airspace monocyte phenotype were confounded with male biological sex, one would expect a positive correlation between these gene sets, as opposed to the observed weak negative correlation with male-upregulated gene set and weak positive correlation with female-upregulated gene set.

**Fig. S4.**
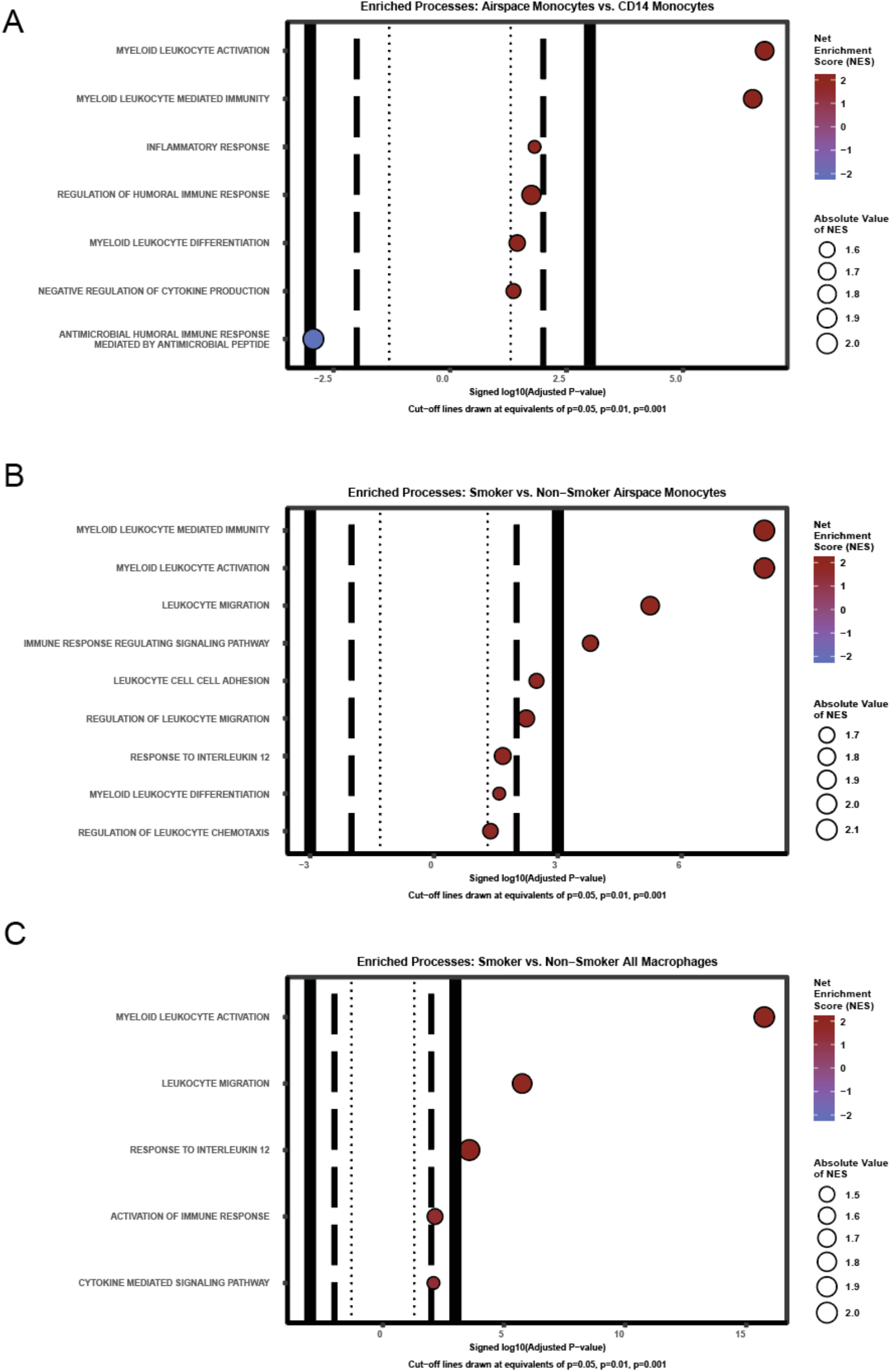
GO Processes Distinguishing Myeloid Populations and Smoking Status. (**A**) GO processes enriched in GSEA analysis of differentially expressed genes between airspace monocytes and CD14 monocyte populations. (**B**) GO processes enriched in GSEA analysis of differentially expressed genes between airspace monocytes derived from smokers vs. airspace monocytes derived from non-smokers. (**C**) GO processes enriched in GSEA analysis of differentially expressed genes between all macrophages derived from smokers vs. all macrophages derived from non-smokers. All presented p-values have undergone Benjamini-Hochberg correction for multiple hypothesis testing.

**Fig. S5.**
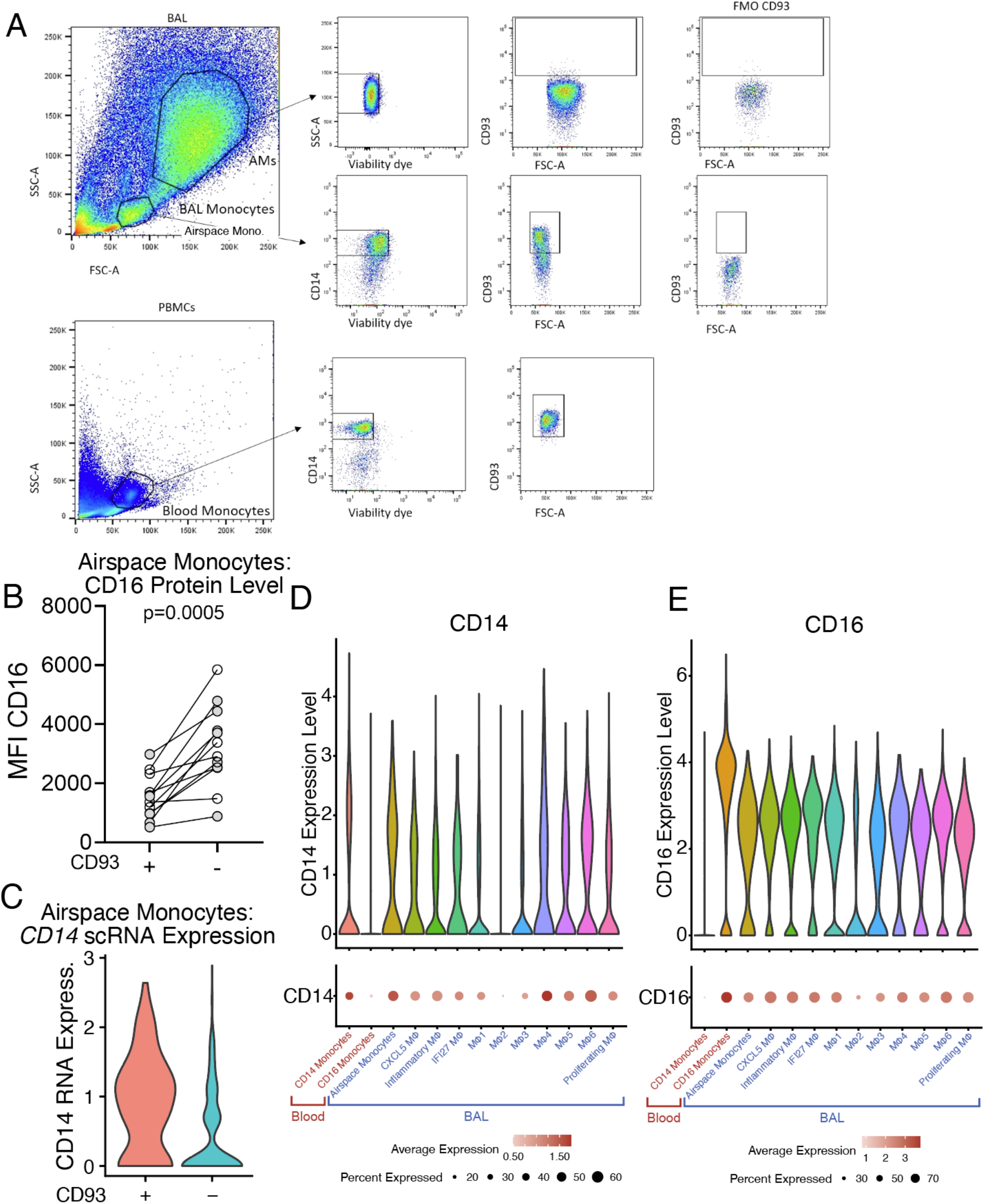
Gating strategy to identify CD93+CD14+ airspace monocytes. **(A)** Alveolar macrophages and airspace monocytes were identified as described in Fig. S1. In addition, CD93 expression of alveolar macrophages, airspace monocytes, and blood monocytes was determined based on FMOs. **(B)** Level of CD16 protein as measured through flow cytometry on airspace monocytes, split by CD93 expression status. **(C)** Level of *CD14* mRNA as measured through scRNA-seq on airspace monocytes, split by CD93 expression status. (**D**) Levels of *CD14* mRNA in scRNA-seq-defined clusters. (**E**) Levels of *CD16* (encoded by *FCGR3A*) mRNA in scRNA-seq-defined clusters.

**Fig. S6.**
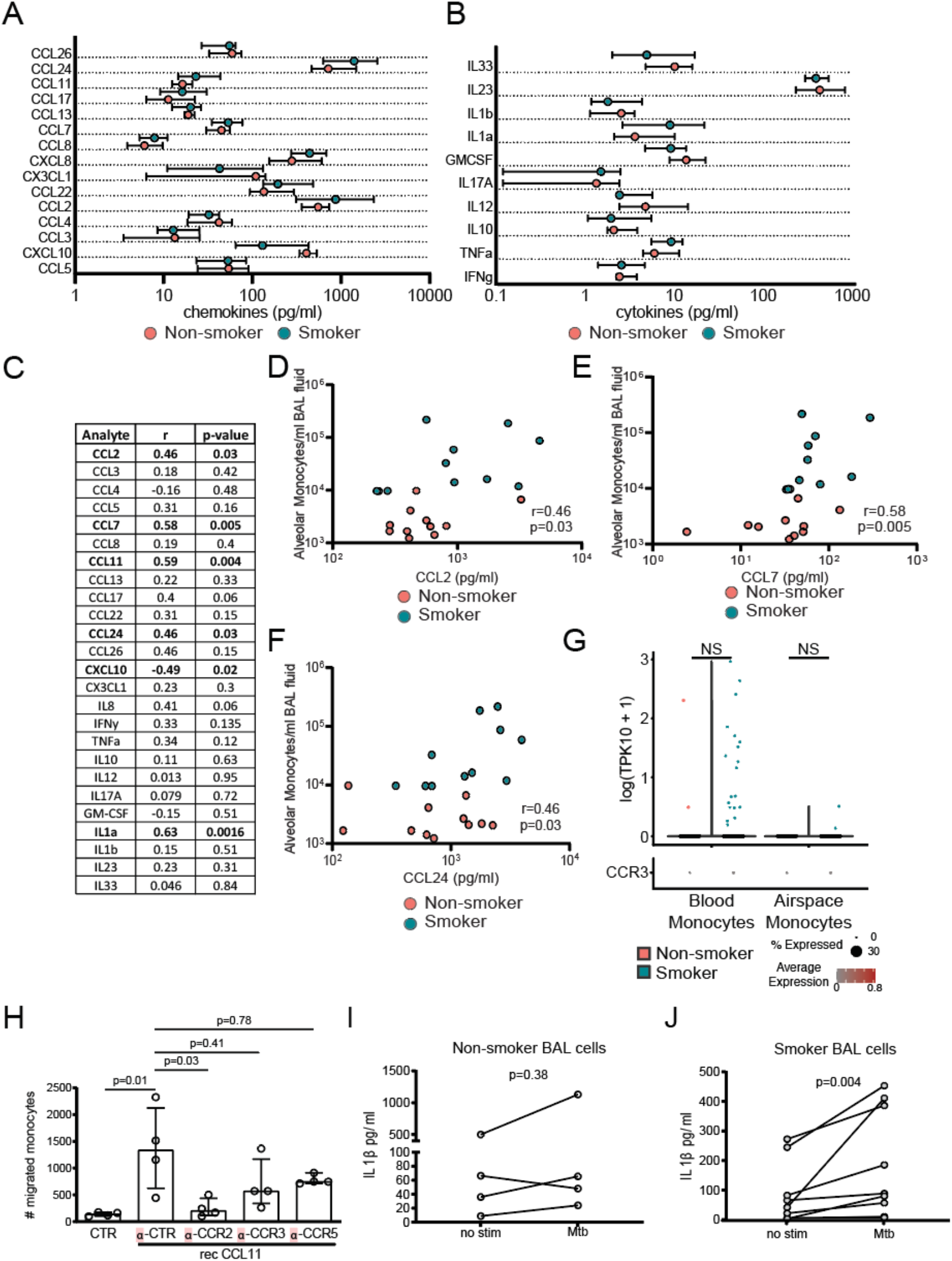
Analysis of soluble factors in BAL from smokers and non-smokers. **(A-C)**. 30x concentrated BAL fluid was analyzed for **(A)** chemokines and **(B)** cytokines using a milliplex magnetic bead panel. **(C)** Correlation of airspace monocyte numbers with BAL chemokine/cytokine concentration with significant correlations in bold and determined by a nonparametric Spearman’s correlation test. **(D-F)** Correlations between airspace monocyte numbers and **(D)** CCL2, **(E)** CCL7, and **(F)** CCL24. **(G)** scRNA-seq expression level of CCR3 in blood and airspace monocytes across non-smokers and smokers. **(H)** Monocyte transwell migration assay towards recombinant CCL11 in the presence of antibodies against CCR2, CCR3, CCR5 or a control antibody. **(I)** IL1-β levels in adherent BAL cells from non-smokers, with or without 24 hours of *Mtb* exposure. **(J)** IL1-β levels in adherent BAL cells from smokers, with or without 24 hours of *Mtb* exposure. P values were determined by a non-parametric Kruskal-Wallis test followed by Dunn’s multiple comparison test. Scatter plots are labelled with median and interquartile range.

**Fig. S7.**
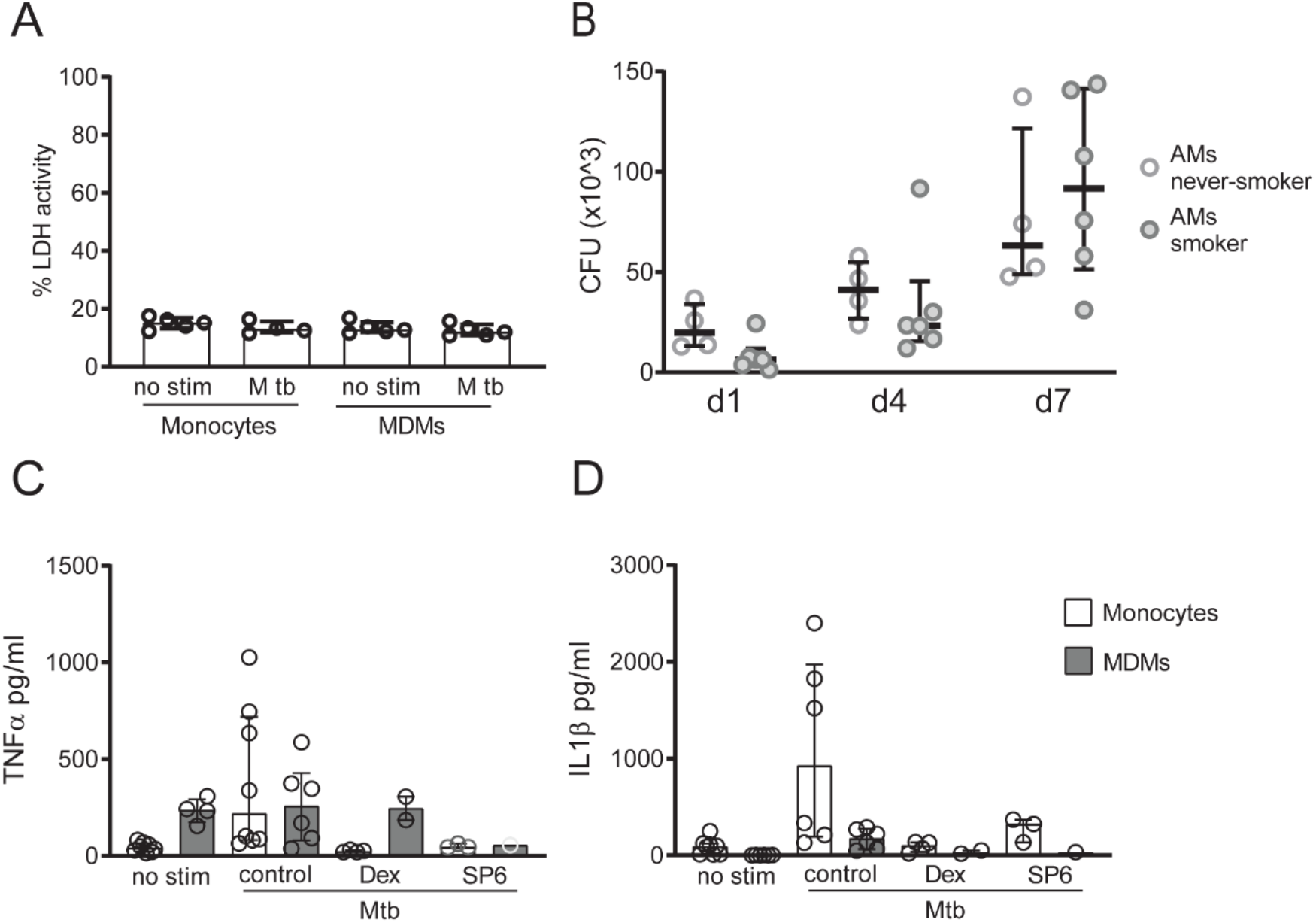
Susceptibility of inflammatory monocytes to intracellular growth of *Mtb*. Monocytes, MDMs or adherent BAL cells were infected with *Mtb* at an MOI of 0.1 for 2h before removing extracellular bacteria and following growth of over time. **(A)** Cell death was determined by LDH activity in the supernatant of monocytes and MDMs exposed to *Mtb* or left untreated for 48h. **(B)** Relative CFU (to day 1) from adherent BAL cells from smokers and non-smokers at day 4 and day 7. **(C)** Extracellular TNF-α and **(D)** IL1-β in cell culture supernatants from monocytes or MDMs exposed to *Mtb* for 24h or left untreated. Where indicated cells were treated with either dexamethasone (40ng/ml) or SP600125 (SP6; 10μM) 30min before infection with *Mtb*. P values were determined by a non-parametric two-way ANOVA followed by Sidak’s multiple comparison test. Scatter plots are labelled with median and interquartile range.

**Table S1.** Characteristics of Human Subjects.

|  | Non-Smokers (*n* = 21) | Smokers (*n* = 13) | Statistics |
| --- | --- | --- | --- |
| male/female | 11/10 | 11/2 | 0.075<sup>a</sup> |
| Age (years) | 35 (24 – 47) | 55 (52 – 58) | **0.0003<sup>b</sup>** |
| Cigarette pack years | N/A | 17 (8-28) | N/A |
| FEV<sub>1</sub> | 4.01 (3.54 – 4.63) | 3.34 (2.65 – 3.78) | **0.01<sup>b</sup>** |
| FEV<sub>1</sub> (% predicted) | 119.5 (105 – 119.5) | 99 (85 – 112) | **0.004<sup>b</sup>** |
| FVC | 5.31 (4.44 – 5.68) | 4.28 (3.56 – 4.79) | **0.04<sup>b</sup>** |
| FVC (% predicted) | 121.5 (115.5 – 130.5) | 102 (84.5 – 116.5) | **0.008<sup>b</sup>** |
| FEV<sub>1</sub>/FVC | 81 (76.75 – 84.5) | 79 (75.5 – 82) | 0.31<sup>b</sup> |

P values were determined by Fisher's exact test<sup>a</sup> or non-parametric Mann-Whitney<sup>b</sup>. Numbers are reported as median with interquartile range. FEV<sub>1</sub>= Forced expiratory volume in 1 second; FVC= Forced vital capacity.

**Table S2:** Antibodies used in this study.

| **Antigen** | **Clone** | **Fluorochrome** | **Company** | **Cat. No.** | **Dilution** |
| --- | --- | --- | --- | --- | --- |
| CD93 | VIMD2 | PE | Biolegend | 336107 | 1:40 |
| CD105 | 43A3 | PE-Cy7 | Biolegend | 323217 | 1:40 |
| CD14 | M5E2 | Alexa 700 | BD | 557923 | 1:50 |
| CD16 | 3G8 | APC-Cy7 | Biolegend | 302018 | 1:50 |
| CD163 | GHI/61 | Texas Red/PE-CF594 | BD | 562670 | 1:100 |
| CD80 | 25F9 | eFluor660 | eBioscience | 50011541 | 1:50 |
| CD14 | M5E2 | Alexa 700 | BD | 557923 | 1:100 |
| CD8 | SK1 | APC | Biolegend | 344722 | 1:100 |
| CD66b | G10F5 | PercP-Cy5.5 | Biolegend | 305108 | 1:200 |
| CD16 | 3G8 | APC-Cy7 | Biolegend | 302018 | 1:50 |
| CD4 | RPTA-4 | BV605 | Biolegend | 300555 | 1:100 |
| CD45 | HI30 | BV570 | Biolegend | 304034 | 1:100 |
| CD3 | UCHT1 | PE-Cy7 | Biolegend | 300420 | 1:100 |

**Table S3:** Results of the Milliplex assay for monocytes and MDMs.

| Analyte | MDMs no stim | MDMs+M tb | p-value | Monocytes no stim | Monocytes + M tb | p-value |
| --- | --- | --- | --- | --- | --- | --- |
| CCL2 | 22415 | 31446 | 0.18 | 2340 | 16953 | 0.63 |
| CCL3 | 181 | 661 | 0.63 | 48 | 39908 | 0.001 |
| CCL4 | 442 | 1016 | 0.44 | 70 | 3264 | 0.0004 |
| CCL7 | 549 | 1047 | 0.44 | 84 | 698 | 0.63 |
| CCL8 | 62 | 106 | 0.44 | 38 | 47 | 0.99 |
| CCL11 | 52 | 58 | 0.36 | 50 | 65 | 0.24 |
| CCL13 | 37 | 53 | 0.74 | 40 | 59 | 0.74 |
| CCL22 | 37805 | 60835 | 0.99 | 765 | 3457 | 0.87 |
| CCL24 | 4571 | 7375 | 0.53 | 33 | 17697 | 0.04 |
| CCL26 | 230 | 77 | 0.99 | 116 | 481 | 0.53 |
| CXCL10 | 141 | 183 | 0.99 | 44 | 35 | 0.99 |
| IL8 | 17263 | 35233 | 0.87 | 22634 | 95543 | 0.04 |
| IL10 | 26 | 34 | 0.99 | 10 | 131 | 0.0024 |
| TNF-α | 440 | 461 | 0.44 | 92 | 3056 | 0.0004 |
| IL12p40 | 3 | 16 | 0.29 | 3 | 16 | 0.29 |
| GM-CSF | 29 | 49 | 0.63 | 31 | 1026 | 0.0051 |
| IL1a | 3 | 5 | 0.99 | 9 | 1695 | 0.03 |
| IL1b | 5 | 6 | 0.99 | 15 | 3448 | 0.03 |
| IL23 | 958 | 1481 | 0.53 | 542 | 1838 | 0.53 |
| IL21 | 21 | 53 | 0.53 | 27 | 86 | 0.08 |

**Table S4:**
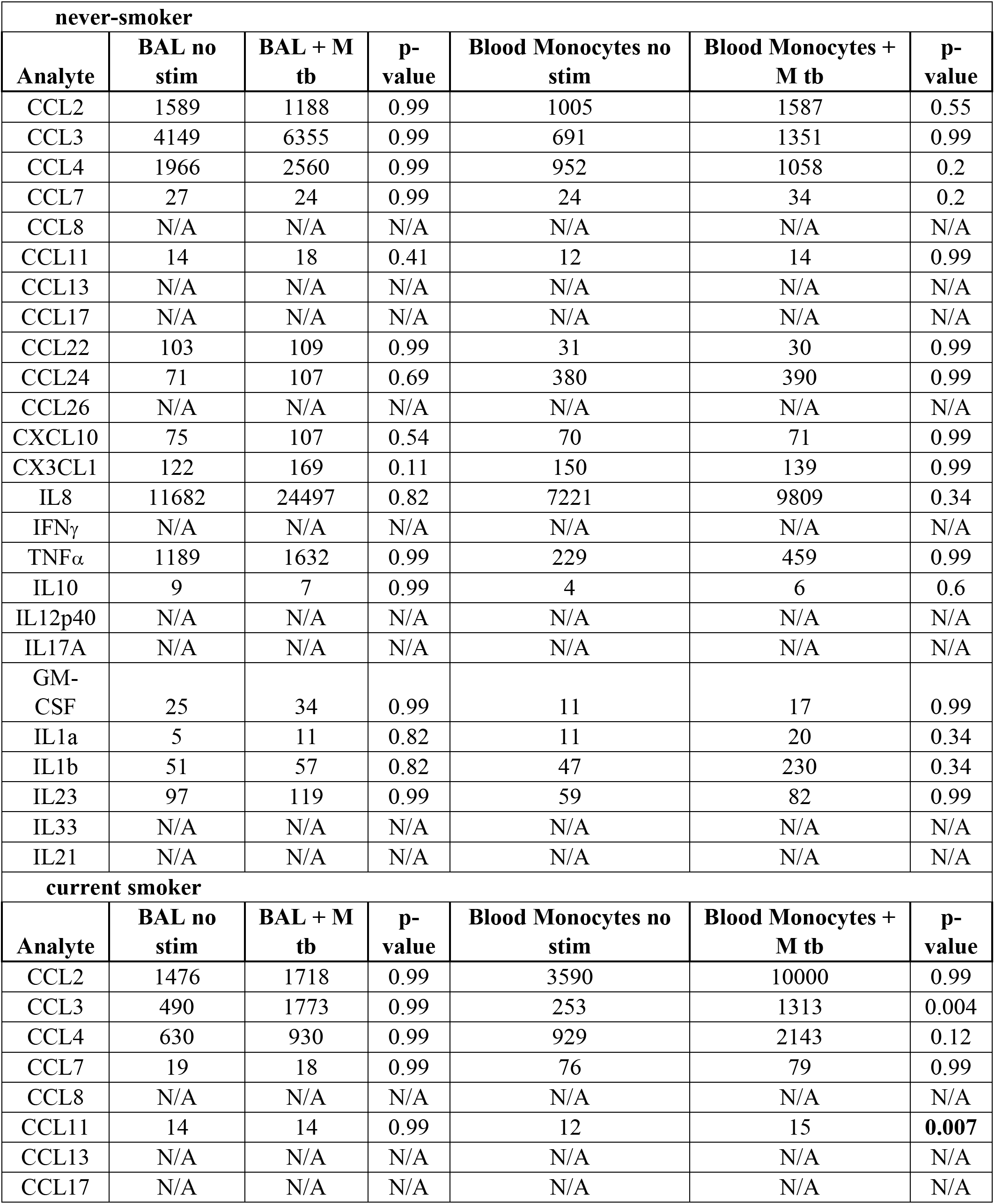
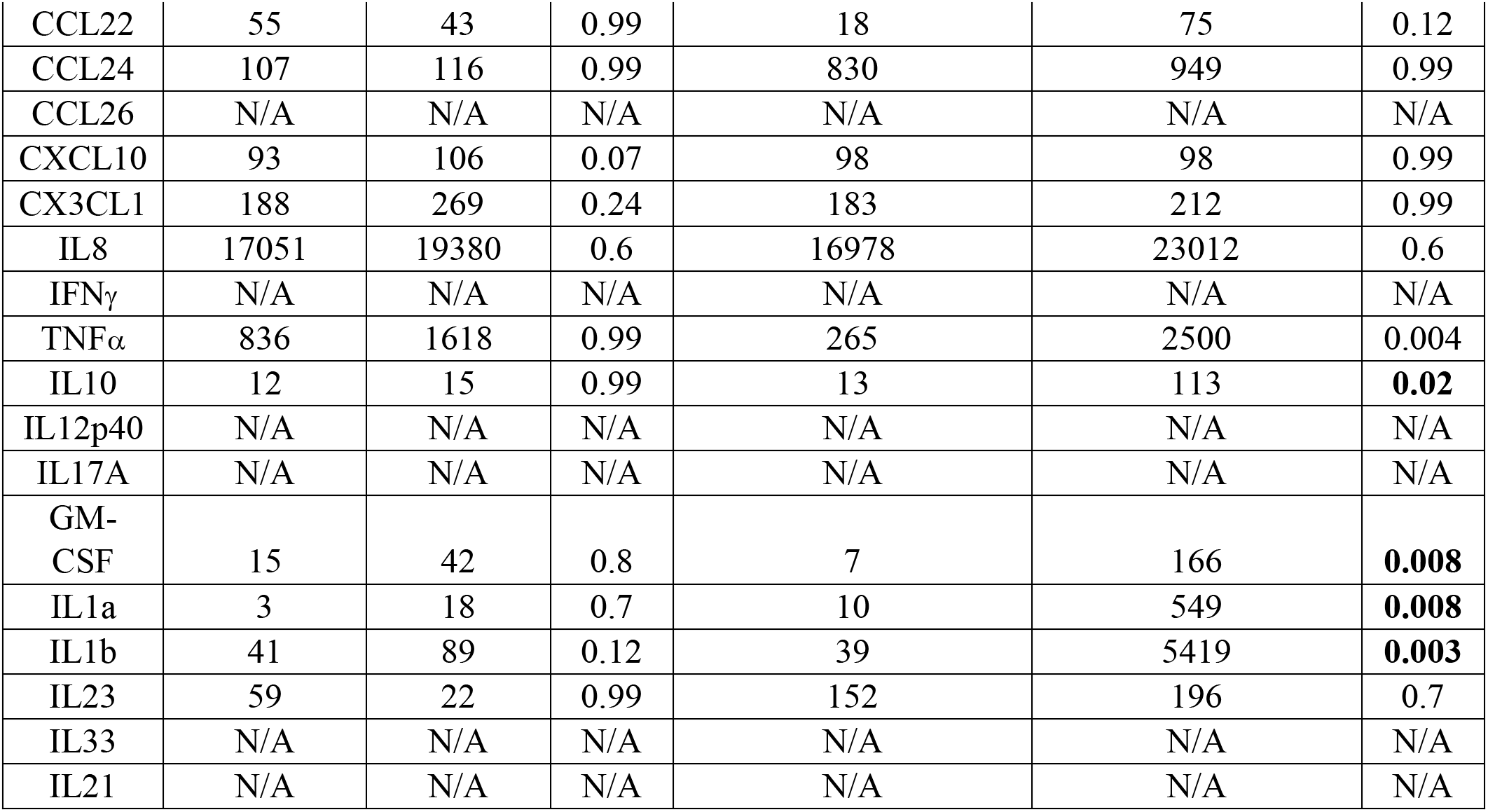
Results Milliplex assay for BAL cells and blood monocytes.

**Table S5:**
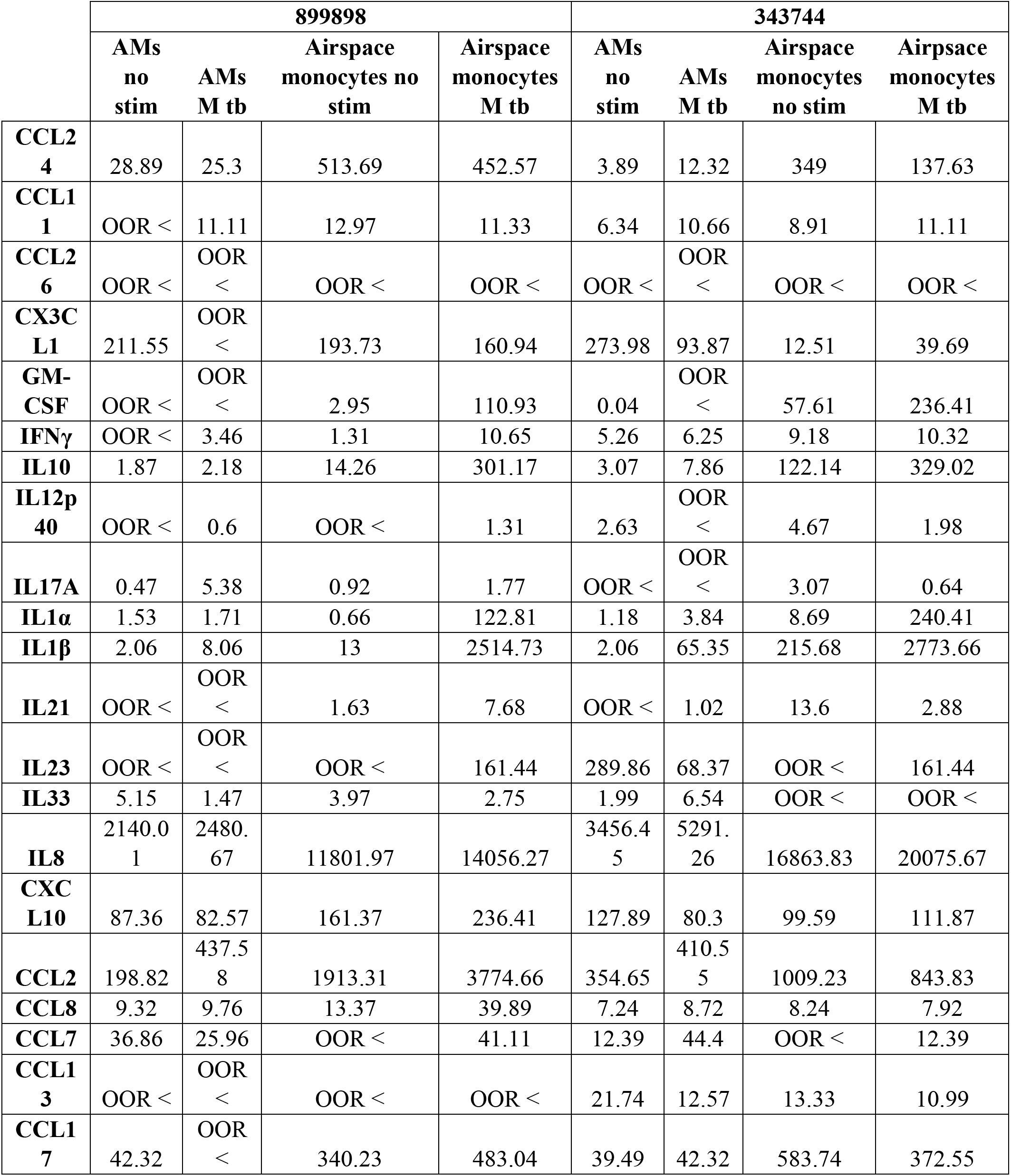
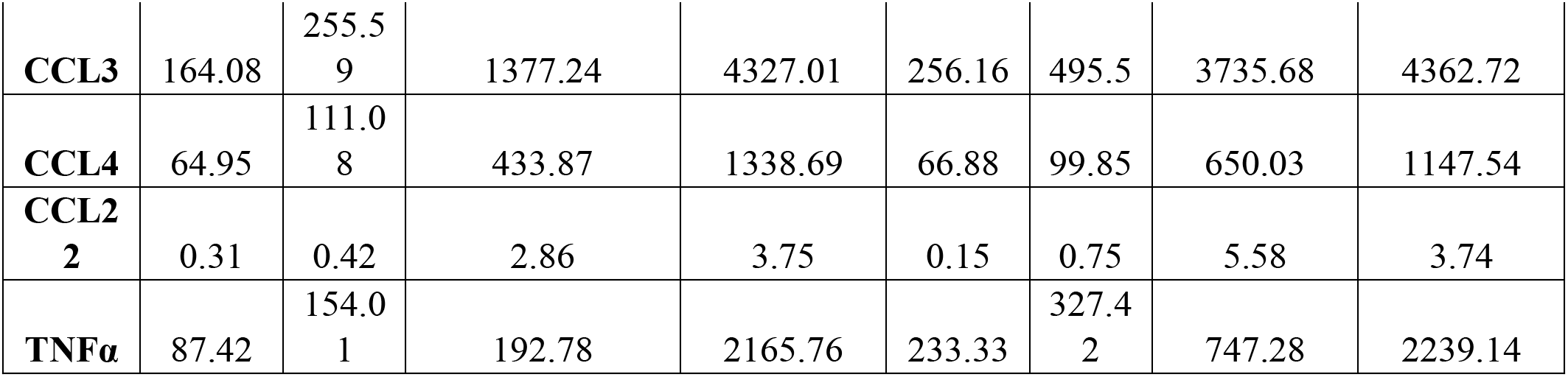
Results of the Milliplex assay for macrophage and monocytes populations isolated from 2 smokers.

